# Protein yield is tunable by synonymous codon changes of translation initiation sites

**DOI:** 10.1101/726752

**Authors:** Bikash K. Bhandari, Chun Shen Lim, Daniela M. Remus, Augustine Chen, Craig van Dolleweerd, Paul P. Gardner

**Affiliations:** Department of Biochemistry, School of Biomedical Sciences, University of Otago, Dunedin, New Zealand; Biomolecular Interaction Centre, University of Canterbury, Christchurch, New Zealand; Callaghan Innovation Protein Science and Engineering, University of Canterbury, Christchurch, New Zealand

**Author notes:** These authors contributed equally (Bikash K. Bhandari, Chun Shen Lim).

## Abstract

Recombinant protein production is a key process in generating proteins of interest in the pharmaceutical industry and biomedical research. However, about 50% of recombinant proteins fail to be expressed in a variety of host cells. To address this problem, we modified up to the first nine codons of messenger RNAs with synonymous substitutions and showed that protein levels can be tuned. These modifications alter the ‘accessibility’ of translation initiation sites. We also reveal the dynamics between accessibility, gene expression, and turnovers using a coarse-grained simulation.

## INTRODUCTION

Recombinant protein expression has numerous applications in biotechnology and biomedical research. Despite extensive refinements in protocols over the past three decades, half of the experiments fail in the expression phase (http://targetdb.rcsb.org/metrics/). Notable problems are the low expression of ‘difficult-to-express’ proteins such as those found in, or associated with, membranes, and the poor growth of the expression hosts, which may relate to toxicity of heterologous proteins (Kimelman et al., 2012) (see (Berlec and Strukelj, 2013; Rosano and Ceccarelli, 2014) for detailed reviews). Despite these issues, mRNA abundance can only explain up to 40% of the variation in protein abundance, due to the complexity of translation and turnover of biomolecules (Abreu et al., 2009; Bernstein et al., 2002; Hanson and Coller, 2018; Lim et al., 2018; Schwanhäusser et al., 2011; Stevens and Brown, 2013; Taniguchi et al., 2010). Furthermore, strong promoters used in expression vectors do not always lead to a desirable level of protein expression because of leaky expression (Rosano and Ceccarelli, 2014).

For *Escherichia coli*, mainstream models that may explain the lower-than-expected correlation between mRNA and protein levels are codon-usage and mRNA structure. Codon analysis is based on the frequency of codon usage in highly expressed proteins using codon adaptation index (CAI) (Sharp and Li, 1987) or tRNA adaptation index (tAI) (Reis and d. Reis, 2004; Sabi and Tuller, 2014), whereas mRNA folding analysis predicts the stability of mRNA secondary structures. Codon usage bias is thought to correlate with tRNA abundance, translation efficiency and protein production (Brule and Grayhack, 2017; Gutman and Hatfield, 1989; Osterman et al., 2020; Reis and d. Reis, 2004; Sabi and Tuller, 2014; Sharp and Li, 1987; Verma et al., 2019) but its usefulness has been questioned (Boël et al., 2016; Cambray et al., 2018; Kudla et al., 2009; Plotkin and Kudla, 2011). More recent studies show stronger support for models based on mRNA folding, in which the stability of RNA structures around the Shine-Dalgarno sequence and translation initiation sites inversely correlates with protein expression (Cambray et al., 2018; de Smit and van Duin, 1990; Dvir et al., 2013; Kudla et al., 2009; Plotkin and Kudla, 2011; Tuller and Zur, 2015). We recently proposed a third model in which the avoidance of inappropriate interactions between mRNAs and non-coding RNAs has a strong effect on protein expression (Umu et al., 2016). The roles of these models in protein expression is an active area of research.

The algorithms for gene optimisation sample synonymous protein-coding sequences using ‘fitness’ models based on CAI, tAI, mRNA folding, and/or G+C content (%) (Chung and Lee, 2012; Raab et al., 2010; Salis et al., 2009; Terai et al., 2016; Villalobos et al., 2006). However, these ‘fitness’ models are usually based on some of the above findings that rely on either endogenous proteins, reporter proteins, or a few heterologous proteins with their synonymous variants. It is unclear whether these features are generalisable to explain the expression of all heterologous proteins. To address this question, we studied multiple large datasets across species in order to extract features that allow us to predict the outcomes of 11,430 experiments of recombinant protein expression in *E. coli*. With this information, we propose how such features can be exploited to fine-tune protein expression at a low cost.

## RESULTS

### Accessibility of translation initiation sites strongly correlates with protein abundance

To identify a better energetic model for mRNA structure that explains protein expression, we examined an *E. coli* expression dataset of green fluorescent protein (GFP) fused in-frame with a library of 96-nt upstream sequences (N=244,000) (Cambray et al., 2018). We removed the redundancy of these 96-nt upstream sequences by clustering on sequence similarity, giving rise to 14,425 representative sequences. We calculated the accessibility (also known as ‘opening energy’ based on unpairing probability) for all the corresponding sub-sequences (see Methods). We examined the correlation between the opening energies and GFP levels. We found that the opening energies of translation initiation sites, in particular from the nucleotide positions −30 to 18 (−30:18), shows the highest correlation with protein abundance (Fig 1A; Spearman’s correlation, R_s_=−0.65, P<2.2×10^−16^). This is stronger than the highest correlation between the minimum free energy −30:30 and protein abundance, which was previously reported as the highest ranked feature (Fig 1A; R_s_=0.51, P<2.2×10^−16^). To account for multiple-testing, the P-values were adjusted using Bonferroni’s correction and reported to machine precision. The datasets used and results are summarised in Supplementary Table S1.

**Fig 1.**
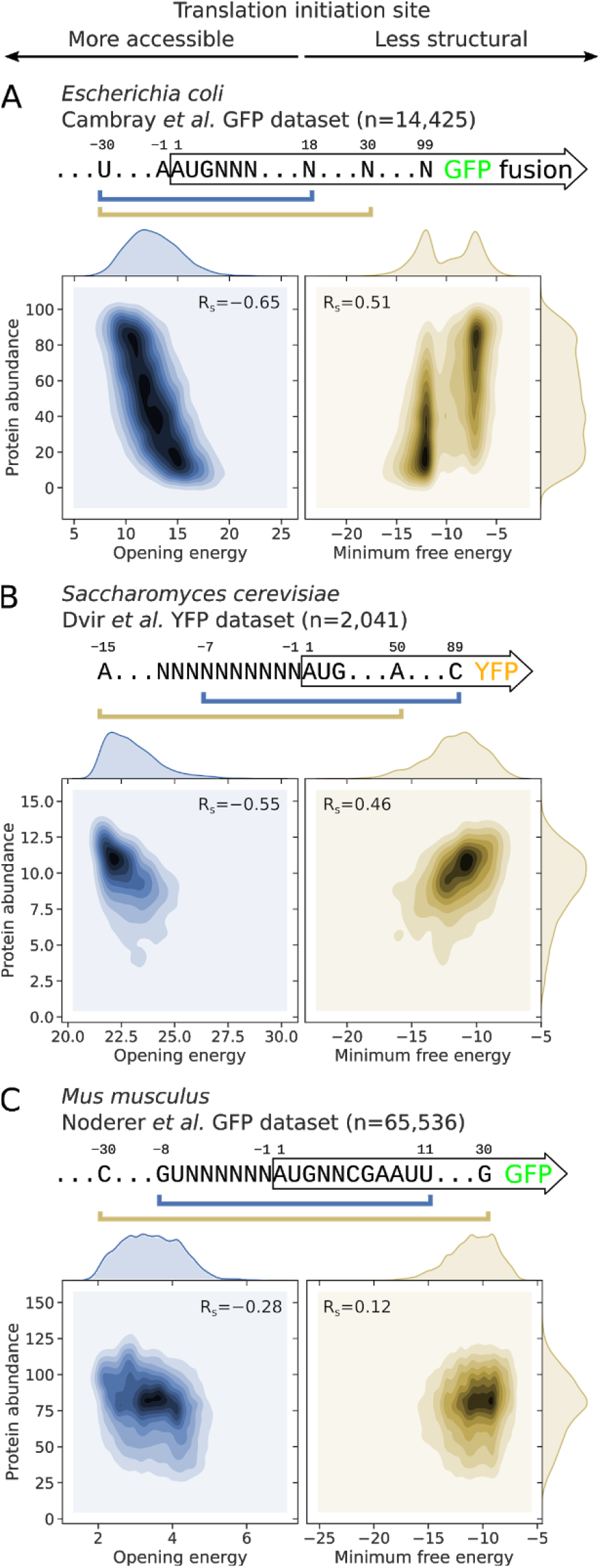
Correlations between the opening energies of translation initiation sites and protein abundance are stronger than that of minimum free energy. **(A)** For *E. coli*, the opening energy at the region −30:18 shows the strongest correlation with protein abundance (also see Fig 2B or Supplementary Fig S1A, sub-sequence l=48 at position i=18). For this analysis, we used a representative GFP expression dataset from Cambray et al. (2018). The reporter library consists of GFP fused in-frame with a library of 96-nt upstream sequences (N=14,425). The minimum free energy −30:30 shown was determined by Cambray et al. (right panel). **(B)** For *S. cerevisiae*, the opening energy −7:89 shows the strongest correlation with protein abundance (also see Supplementary Fig S1B, sub-sequence l=96 at position i= 89). For this analysis, we used the YFP expression dataset from Dvir et al. (2013). The YFP reporter library consists of 2,041 random decameric nucleotides inserted at the upstream of YFP start codon. The minimum free energy −15:50 was previously shown to correlate the best with protein abundance (right panel). **(C)** For *M. musculus*, the opening energy −8:11 shows the strongest correlation with protein abundance (also see Supplementary Fig S1C, sub-sequence l=19 at position i=11). For this analysis, we used the GFP expression dataset from Noderer et al. (2014). The GFP reporter library consists of 65,536 random hexameric and dimeric nucleotides inserted at the upstream and downstream of GFP start codon, respectively. The minimum free energy −30:30 was shown (right panel). See also Supplementary Table S1. R_s_, Spearman’s rho. Bonferroni adjusted P-values are below machine’s underflow level for the correlations between opening energies and protein abundances shown in the left panels.

We repeated the analysis for a dataset of yellow fluorescent protein (YFP) expression in *Saccharomyces cerevisiae (Dvir et al*., *2013)*. This dataset corresponds to a library of 5′UTR variants, in which the 10-nt sequences preceding the YFP translation initiation site were randomly substituted (N=2,041). In this case, the opening energy −7:89 showed a stronger correlation with protein abundance than that of the minimum free energy −15:50 reported previously (Fig 1B; R_s_=−0.55 versus 0.46).

To examine the usefulness of accessibility in complex eukaryotes, we analysed a dataset of GFP expression in *Mus musculus (Noderer et al*., *2014)*. The reporter library was originally designed to measure the strength of translation initiation sequence context, in which the 6- and 2-nt sequences upstream and downstream of the GFP translation initiation site were randomly substituted, respectively (N=65,536). Here the opening energy −8:11 showed a maximum correlation with expressed proteins, which again, is stronger than that of the minimum free energy −30:30 (Fig 1C; R_s_=−0.28 versus 0.12).

Taken together, our findings suggest that the accessibility of translation initiation sites strongly correlates with protein abundance across species. Interestingly, our findings also suggest that the Shine-Dalgarno sequence (Shine and Dalgarno, 1974) at −13:−8 should be accessible to recruit ribosomes.

### Accessibility predicts the outcome of recombinant protein expression

We investigated how accessibility performs in the real world in prediction of recombinant protein expression. For this purpose, we analysed 11,430 expression experiments in *E. coli* from the ‘Protein Structure Initiative:Biology’ (PSI:Biology) (Acton et al., 2005; Chen et al., 2004; Seiler et al., 2014). These PSI:Biology targets were expressed using the pET21_NESG expression vector that harbours the *T7lac* inducible promoter and a C-terminal His tag (Acton et al., 2005).

We split the experimental results of the PSI:Biology targets into protein expression ‘success’ and ‘failure’ groups (N=8,780 and 2,650, respectively; see Supplementary Fig S2). These PSI:Biology targets span more than 189 species and the failures are representative of various problems in heterologous protein expression. Only 1.6% of the targets were *E. coli* proteins, which is negligible (N=179; see Supplementary Fig S2).

We calculated the opening energies for all possible sub-sequences of the PSI:Biology targets as above (Fig 2, positions relative to initiation codons). For each sub-sequence region, we used the opening energies to predict the expression outcomes and computed the prediction accuracy using the area under the receiver operating characteristic curve (AUC; see Fig 2C). A closer look into the correlations between opening energies and expression outcomes, and AUC scores calculated for the sub-sequence regions reveals a strong accessibility signal of translation initiation sites (Fig 2B&C, Cambray’s GFP and PSI:Biology datasets, respectively). We matched the correlations and AUC scores by sub-sequence regions and confirmed that sub-sequence regions that have strong correlations are likely to have high AUC scores (Fig 2D). In contrast, the sub-sequence regions that have zero correlations are not useful for predicting the expression outcomes (AUC approximately 0.5).

**Fig 2.**
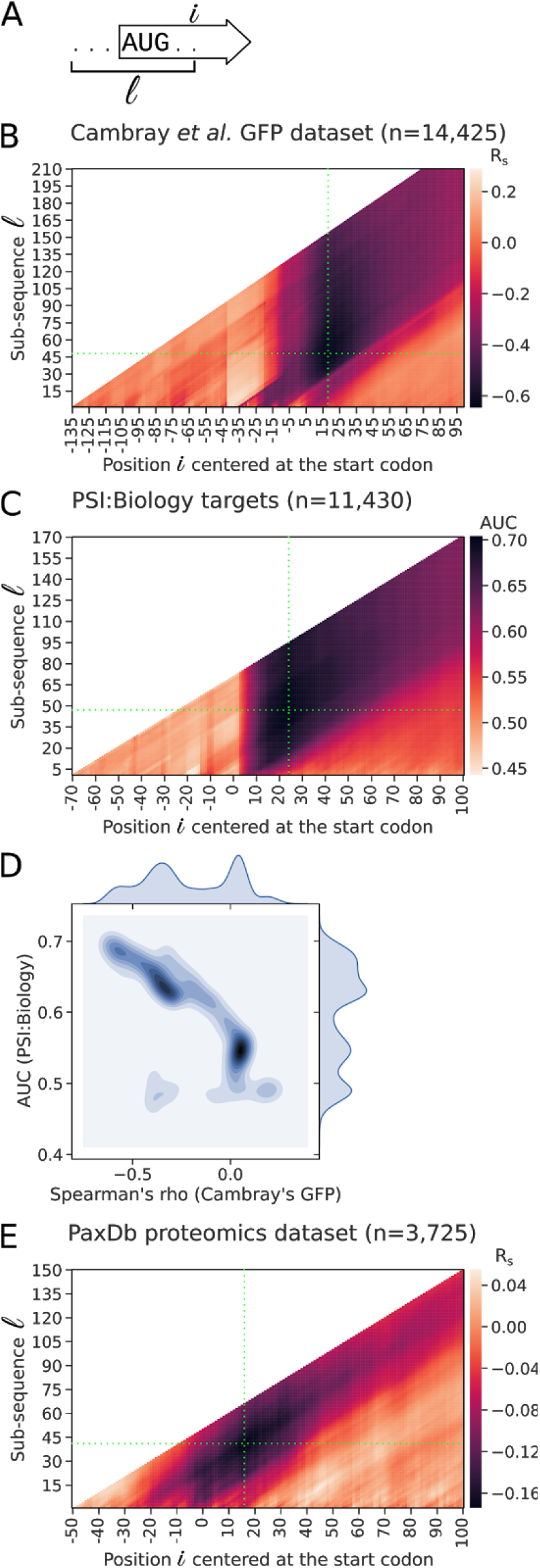
Opening energies of regions surrounding the Shine-Dalgarno and start codons are predictive of protein expression in *E. coli*. **(A)** Schematic representation of a transcript sub-sequence l at position i for the calculation of opening energy. For example, sub-sequence l=10 at position i=10 corresponds to the region 1:10. **(B)** Correlation between the opening energies for the sub-sequences of GFP transcripts and protein abundance. The opening energy at the region −30 to 18 nt (sub-sequence l=48 at position i=18, green crosshair) shows the strongest correlation with protein abundance [R_s_=−0.65; N=14,425, GFP expression dataset of Cambray et al. (2018)]. For this dataset, the reporter plasmid used is pGC4750, in which the promoter and ribosomal binding site are oFAB1806 inducible promoter and oFAB1173/BCD7, respectively. **(C)** Prediction accuracy of the expression outcomes of the PSI:Biology targets using opening energy (N=11,430). The opening energy at the region −23:24 (sub-sequence l=47 at position i=24, green crosshair) shows the highest prediction accuracy score (AUC=0.70). For this dataset, the expression vector used is pET21_NESG, in which the promoter and fusion tag are T7lac and C-terminal His tag, respectively. **(D)** Comparison between the correlations and AUC scores by sub-sequence region taken from the above analyses. The sub-sequence regions that have strong correlations are likely to have high AUC scores, whereas the sub-sequence regions that have no correlations are likely not useful in prediction of the expression outcomes. **(E)** Correlation between the opening energies for the sub-sequences of *E. coli* transcripts and protein abundance. The transcripts used for this analysis are protein-coding sequences concatenated with 50 and 10 nt located upstream and downstream, respectively. The opening energy at the region −25:16 (sub-sequence l=41 at position i=16, green crosshair) shows the strongest correlation with protein abundance (R_s_=−0.17; N=3,725, PaxDb integrated proteomics dataset). See also Supplementary Table S1. R_s_, Spearman’s rho.

We then asked how accessibility manifests in the endogenous mRNAs of *E. coli*, for which we studied a proteomics dataset of 3,725 proteins available from PaxDb (Wang et al., 2015). As expected, we observed a similar accessibility signal, with the region −25:16 correlated the most with protein abundance (Fig 2E). However, the correlation was rather low (R=−0.17, P<2.2×10^−16^), which may reflect the limitation of mass spectrometry to detect lower abundances (Nilsson et al., 2010; Tabb et al., 2009). Furthermore, the endogenous promoters have variable strength, which gives rise to a broad range of mRNA and protein levels (Delvigne et al., 2017; Deuschle et al., 1986). Taken together, our results show that the accessibility signal of translation initiation sites is very consistent across various datasets analysed (Supplementary Fig S1 and Fig 2).

### Accessibility outperforms other features in prediction of recombinant protein expression

To choose an accessibility region for subsequent analyses, we selected the top 200 regions from the above correlation analysis on Cambray’s dataset (Fig 2B) and used random forest to rank their Gini importance scores in prediction of the outcomes of the PSI:Biology targets. The region −24:24 was ranked first, which is nearly identical to the region −23:24 with the top AUC score (Fig 2C, AUC=0.70). We therefore used the opening energy at the region −24:24 in subsequent analyses.

We asked how the other features perform compared to accessibility in prediction of heterologous protein expression, for which we analysed the same PSI:Biology dataset. We first calculated the minimum free energy and avoidance at the regions −30:30 and 1:30, respectively. These are the local features associated with translation initiation rate. We also calculated CAI (Sharp and Li, 1987), tAI (Tuller et al., 2010), codon context (CC) (Ang et al., 2016), G+C content, and Iχnos scores (Tunney et al., 2018). CC is similar to CAI except it takes codon-pairs into account, whereas the Iχnos scores are translation elongation rates predicted using a neural network model trained with ribosome profiling data (Supplementary Fig S3). These are the global features associated with translation elongation rate. We built a random forest model to rank the Gini importance scores of these local and global features. The local features ranked higher than the global features (Fig 3A). We then calculated and compared the prediction accuracy of these features. The AUC scores for the local features were 0.70, 0.67 and 0.62 for the opening energy, minimum free energy and avoidance, respectively, whereas the global features were 0.58, 0.57, 0.54, 0.54 and 0.51 for Iχnos, G+C content, CAI, CC and tAI, respectively (Fig 3B). The local features outperform the global features, suggesting that effects on translation initiation are a major predictor of the outcome of heterologous protein expression. We further examined the local G+C contents corresponding to the local features (Supplementary Fig S4). The G+C contents in the regions −24:24 and −30:30 weakly correlate with opening energy and minimum free energy, respectively. The AUC scores for these local G+C contents are also lower than the corresponding local features, suggesting that these local G+C contents are not good proxies for the corresponding local features. Overall, our findings support previous reports that the effects on translation initiation are rate-limiting (Kudla et al., 2009; Tuller and Zur, 2015) which, interestingly, correlate with the binary outcome of recombinant protein expression (Fig 3C). Importantly, accessibility outperformed all other features.

**Fig 3.**
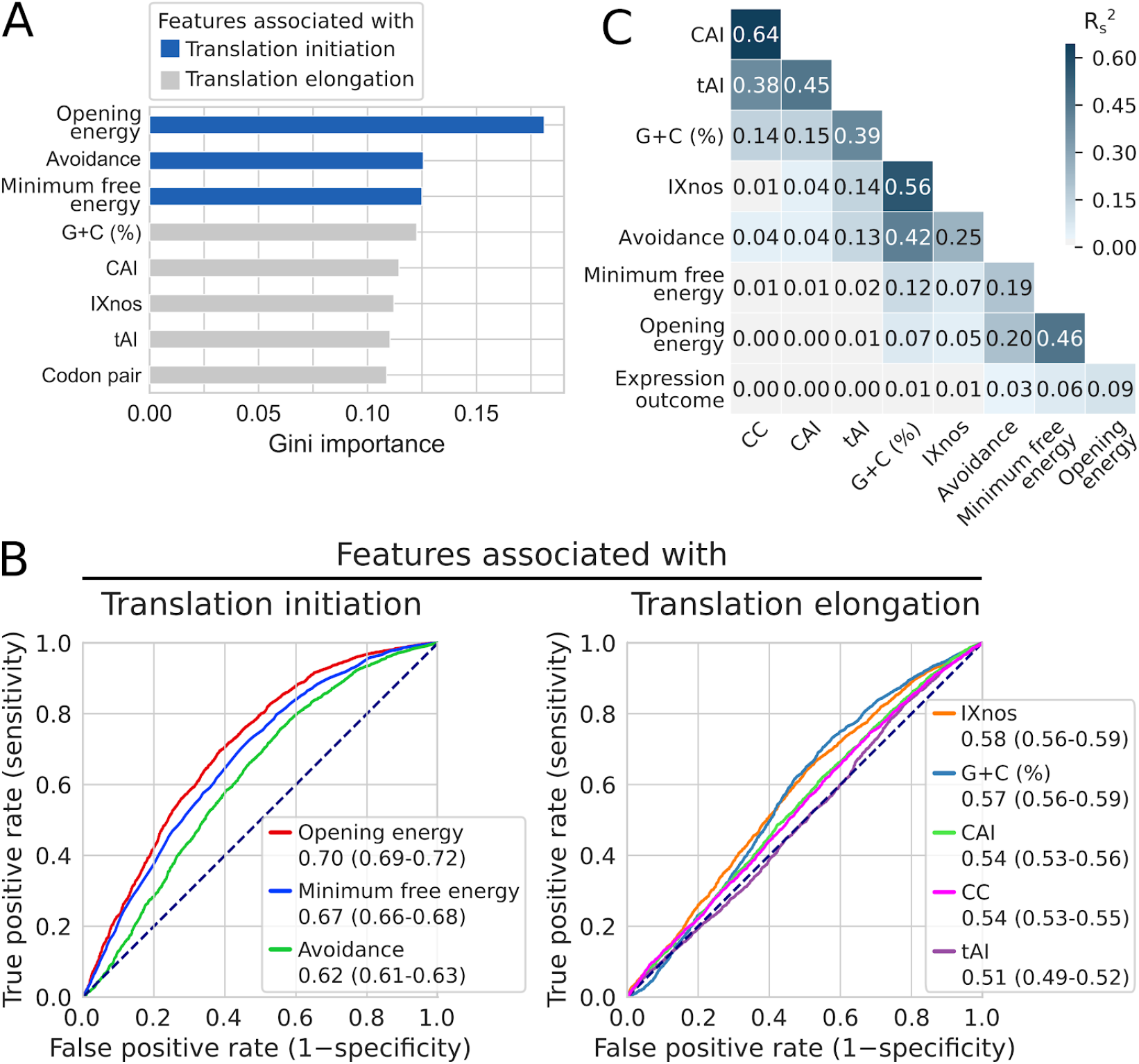
Accessibility of translation initiation sites is the strongest predictor of heterologous protein expression in *E. coli*. **(A)** mRNA features ranked by Gini importance for random forest classification of the expression outcomes of the PSI:Biology targets (N=8,780 and 2,650, ‘success’ and ‘failure’ groups, respectively). The features associated with translation initiation rate (purple; opening energy −24:24, minimum free energy −30:30, and mRNA:ncRNA avoidance 1:30) have higher scores than the feature associated with translation elongation rate [blue; tRNA adaptation index (tAI), codon context (CC), codon adaptation index (CAI), G+C content (%), and Iχnos]. The Iχnos scores are translation elongation rates predicted using a neural network model trained with ribosome profiling data (Supplementary Fig S3). **(B)** ROC analysis shows that accessibility (opening energy −24:24) has the highest classification accuracy. The AUC scores with 95% confidence intervals are shown. See also Supplementary Table S1. **(C)** Accessibility (opening energy −24:24) is the best feature in explaining the expression outcomes. Relationships between the features and expression outcomes represented as squared Spearman’s rho (R_s_^2^).

To identify a good opening energy threshold, we calculated positive likelihood ratios for different opening energy thresholds using the cumulative frequencies of true negative, false negative, true positive and false positive derived from the above receiver operating characteristic (ROC) analysis (Supplementary Fig S5, top panel). Meanwhile, we calculated the 95% confidence intervals of these positive likelihood ratios using 10,000 bootstrap replicates. We reasoned that there is an upper and lower bound on translation initiation rate, therefore the relationship between translation initiation rate and accessibility is likely to follow a sigmoidal pattern. We fit the positive likelihood ratios into a four-parametric logistic regression model (Supplementary Fig S5). As a result, we are 95% confident that an opening energy of 10 kcal/mol or below at the region −24:24 is about two times more likely to belong to the sequences which are successfully expressed than those that failed.

### Accessibility can be improved using a simulated annealing algorithm

The above results suggest that accessibility can, in part, explain the low expression problem of heterologous protein expression. Therefore, we sought to exploit this idea for optimising gene expression. We developed a simulated annealing algorithm to maximise the accessibility at the region −24:24 using synonymous codon substitution (see Methods). Previous studies have found that full-length synonymous codon-substituted transgenes may produce unexpected results, such as a reduction in mRNA abundance, RNA toxicity, and/or protein misfolding (Ben-Yehezkel et al., 2015; Mittal et al., 2018; Tunney et al., 2018; Umu et al., 2016). Therefore, we sought to determine the minimum number of codons required for synonymous substitutions in order to achieve near-optimum accessibility. For this purpose, we used the PSI:Biology targets that failed to be expressed. We applied our simulated annealing algorithm such that synonymous substitutions can happen at any codon of the sequences except the start and stop codons, although the changes may not necessarily happen to all codons due to the stochastic nature of our optimisation algorithm (see Methods). Next, we constrained synonymous codon substitution to the first 14 codons and applied the same procedure (Supplementary Fig S6A). Therefore, the changes may only occur at any or all of the first 14 codons. We repeated the same procedure for the first nine and also the first four codons. Thus a total of four series of codon-substituted sequences were generated. We then compared the distributions of opening energy −24:24 for these series using the Kolmogorov-Smirnov statistic (D_KS_; see Supplementary Fig S6B). The distance between the distributions of the nine and full-length codon-substituted series was significantly different yet sufficiently close (D_KS_=0.087, ×P=3.3 10^−8^), suggesting that optimisation of the first nine codons is sufficient in most cases to achieve an optimum accessibility of translation initiation sites. We named our software <u>T</u>ranslation Initiation coding region des<u>igner</u> (TIsigner), which by default, allows synonymous substitutions in the first nine codons.

We asked to what extent the existing gene optimisation tools modify the accessibility of translation initiation sites. For this purpose, we first submitted the PSI:Biology targets that failed to be expressed to the ExpOptimizer web server from NovoPro Bioscience (see Methods). We also optimised the PSI:Biology targets using the standalone version of Codon Optimisation OnLine (COOL) (Chung and Lee, 2012). We found that both tools increase accessibility indirectly even though their algorithms are not specifically designed to do so. In fact, a purely random synonymous codon substitution on these PSI:Biology targets using our own script resulted in similar increases in accessibility (Supplementary Fig S6C). These results may explain some indirect benefits from the existing gene optimisation tools (i.e. any change from suboptimal is likely to be an improvement, see below).

### Low protein yields can be improved by synonymous codon changes in the vicinity of translation initiation sites

To demonstrate that heterologous protein expression is tunable with minimum effort, we designed and tested a series of GFP reporter gene constructs. We tested 29 plasmids harbouring GFP reporter genes with synonymous changes within the first nine codons (opening energies of 5.56-21.68 kcal/mol; Supplementary Table S2 and Supplementary Methods). GFP expression is controlled by an IPTG inducible *T7lac* promoter. In addition, all plasmids harbour a second reporter gene, i.e. mScarlet-I, which is controlled by the constitutive promoter from the *nptII* gene for aminoglycoside-3’-O-phosphotransferase of *E. coli* transposon Tn5 (Bindels et al., 2017; Schlechter et al., 2018). mScarlet-I expression was measured to correct for plasmid copy number and as a proxy for bacterial growth (Schlechter et al., 2020). As expected, the GFP level significantly correlates with accessibility (i.e., anti-correlates with opening energy, R _s_ =−0.53, P=3.4×10^−3^; Fig 6A). Curiously, we observed a diminishing return with opening energies lower than that of the wild-type sequence (11.68 kcal/mol). To investigate this, we simulated a protein production experiment by modelling cell growth, transcription, translation, and turnovers (see Methods). We assumed that opening energies of 12 kcal/mol or below is favourable in this model, based on our analysis of 8,780 PSI:Biology ‘success’ group (Supplementary Fig S6). Interestingly, our *in silico* coarse-grained model shows a similar protein production trend as the actual experiment (Fig 6B).

We then tested this finding using the luciferase reporter of *Renilla reniformis* (RLuc). Similarly, we designed a series of RLuc variants, but with opening energies below that of the wild-type sequence (5.77-10.38 kcal/mol; Fig 6C and Supplementary Table S2). In addition, we tested commercially designed sequences, in which sequence optimisations were performed in full-length rather than the first 9 codons. We observed that TIsigner (9.9 kcal/mol) and commercially optimised luciferase reporter genes produced significantly higher luminescence than the wild-type (Fig 6C), although RLuc is poorly soluble in the *E. coli* host (Supplementary Fig S8). We also found that the levels of wild-type luciferase and many variants with lower opening energies (5-7 kcal/mol) were not significantly different.

As both wild-type GFP and RLuc genes are strongly expressed in *E. coli*, we asked whether poorly expressed proteins can be improved by increasing accessibility of translation initiation sites. We performed densitometric analysis of previously published Western blots, which include the results of a cell-free expression system using constructs harbouring a wild-type antibody fragment or archaebacterial dioxygenase and its synonymous variants (within the first six codons) (Voges et al., 2004). Indeed, variants with opening energies lower than the wild-type sequences were expressed at higher levels (Fig 6D).

## DISCUSSION

Our findings show that the accessibility of translation initiation sites is the strongest predictor of heterologous protein expression in *E. coli*. Whereas previous studies have largely used minimum free energy models to define the accessibility of a region of interest (Bhattacharyya et al., 2018; Nieuwkoop et al., 2019; Pelletier and Sonenberg, 1987; Salis et al., 2009; Voges et al., 2004). However, Terai and Asai (2020) and ourselves have independently discovered that the opening energy is a better choice for modelling accessibility (Bhandari et al., 2019; Terai and Asai, 2020) (see Fig 1A for example). Opening energy is an ensemble average energy that accounts for suboptimal RNA structures that are not reported by minimum free energy models by default (Bernhart et al., 2011; Mückstein et al., 2006). Currently, the modelling of accessibility using opening energy is largely used for the prediction of RNA-RNA intermolecular interactions, for example, as implemented in RNAup and IntaRNA (Lorenz et al., 2011; Mann et al., 2017). Our study has shown that this approach can be used to identify the key accessibility regions that are consistent across multiple large expression datasets. We have implemented our findings in TIsigner web server, which currently supports recombinant protein expression in *E. coli* and *S. cerevisiae* (optimisation regions −24:24 and −7:89, respectively; see Fig 1). An independent yet similar implementation is available in XenoExpressO web server with the purpose of optimising protein expression for an *E. coli* cell-free system (Zayni et al., 2018). The authors showed that an increase in accessibility of a 30 bp region from the Shine-Dalgarno sequence enhances the expression level of human voltage dependent anion channel, which further supports our findings.

The strengths of our approaches are five-fold. Firstly, the likelihood of success or failure can be assessed prior to running an experiment. Users can compare the opening energies calculated for the input and optimised sequences and the distributions of the ‘success’ and ‘failure’ of the PSI:Biology targets. We also introduced a scoring scheme to score the input and optimised sequences based upon how likely they are to be expressed (Supplementary Fig S5; also see Methods). Secondly, optimised sequences can have up to the first nine codons substituted (by default), meaning that gene optimisation using a standard PCR cloning method is feasible. For cloning, we propose a nested PCR approach, in which the final PCR reaction utilises a forward primer designed according to the optimised sequence (Sambrook and Russell, 2001) (Supplementary Fig S6D). Thirdly, the cost of gene optimisation can be reduced dramatically as gene synthesis is replaced with PCR using our approach. This enables high-throughput protein expression screening using the optimised sequences, generated at a low cost. Fourthly, tunable expression is possible, i.e. high, intermediate or even low expression 5′ codon sequences can be designed, allowing for more control over heterologous protein production, as demonstrated by our experiments (Fig 4). Finally, our fast, lightweight, coarse-grained simulation approach has opened up new avenues to study several aspects of gene expression, such as transcription, translation, cellular growth, and turnovers, which give good proxies to how cellular systems behave.

**Fig 4.**
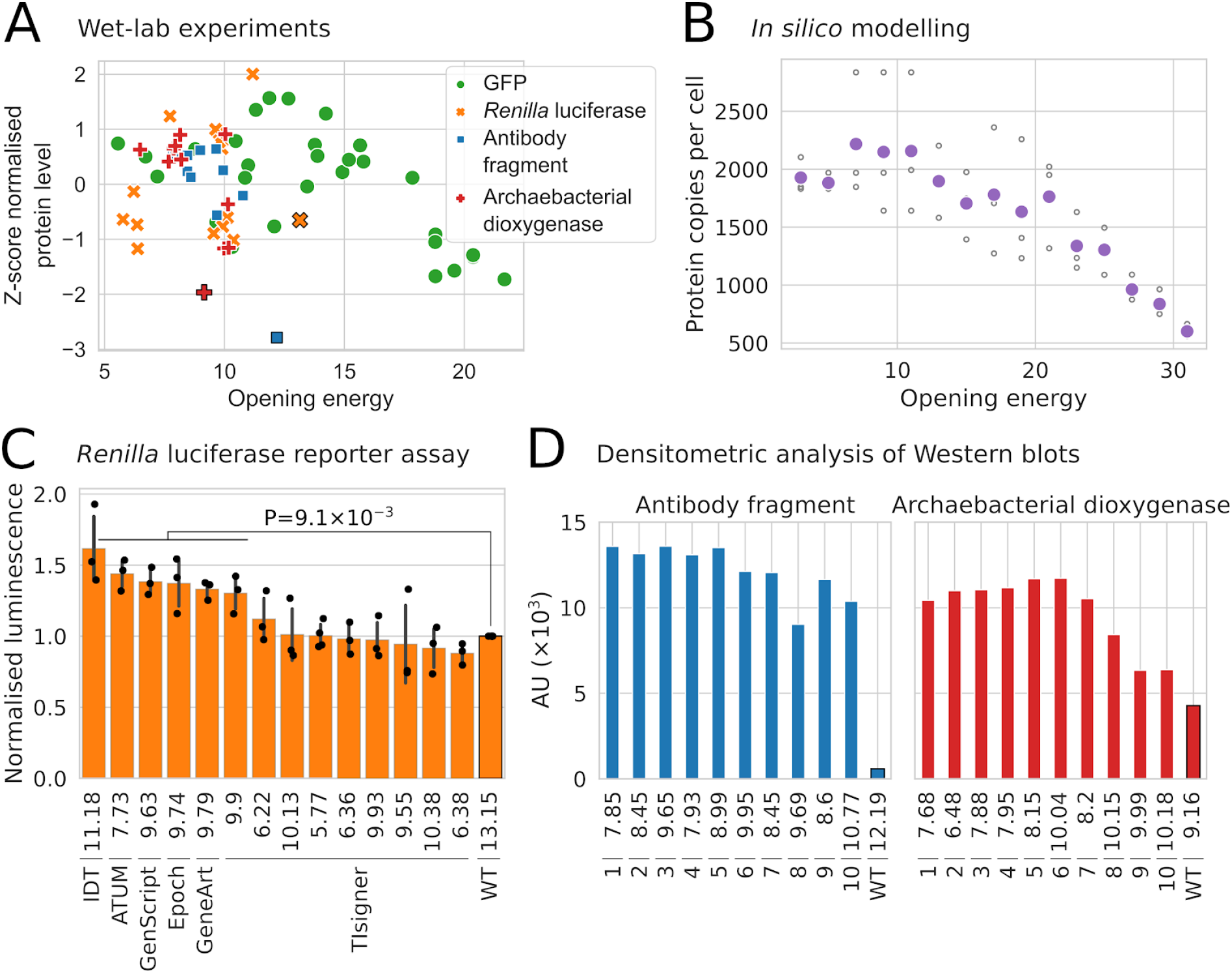
The yields of heterologous protein productions are tunable by synonymous codon changes in the first nine codons. **(A)** GFP level strongly correlates with accessibility, i.e., anti-correlates with opening energy (R_s_=−0.53, P=3.4×10^−3^; N=29). The protein levels of GFP, *Renilla* luciferase (RLuc), an antibody fragment and an archaebacterial dioxygenase were transformed using z-score method. The GFP and RLuc levels were derived from the average values of at least two and three independent biological replicates, respectively. Black outlines denote wild-type sequences. **(B)** Coarse-grained simulation of a protein production experiment by modelling cell growth, transcription, translation, and turnovers, given that translation initiation sites with opening energies less than or equal to 12 kcal/mol is optimum. The *in silico* model shows a similar trend of protein production as the wet-lab experimental results. Unfilled and filled (purple) circles denote the *in silico* replicates and their corresponding average values, respectively (R_s_=−0.75, P=2.8×10^−9^). **(C)** The expression of RLuc can be improved, despite its poor solubility in *E. coli* (Supplementary Fig S8). Opening energies are shown next to labels. The luciferase activities of commercially designed RLuc reporter genes (full-length sequence optimisation) and TIsigner (9.9 kcal/mol) are significantly higher than the wild-type luciferase (Mann-Whitney U tests, P=9.1×10^−3^). No significant differences were observed between the commercial designs and TIsigner (9.9 kcal/mol). Error bars denote standard deviation of three independent biological replicates. **(D)** Densitometric analysis of previously published Western blots shows that the yields of an antibody fragment and an archaebacterial dioxygenase can be improved by synonymous codon changes within the first six codons (Voges et al., 2004). A RTS *E. coli* cell-free expression system was used. The processed data are available Supplementary Table S2. AU, arbitrary unit; R_s_, Spearman’s rho; WT, wild-type.

## MATERIALS AND METHODS

### Sequence features analysis

Datasets used in this study are listed in Supplementary Table S1. Representative sequences were chosen using CD-HIT-EST (Fu et al., 2012; Li and Godzik, 2006). Minimum free energies, opening energies and avoidance were calculated using RNAfold, RNAplfold and RNAup from ViennaRNA package (version 2.4.11), respectively (Bernhart et al., n.d., 2011; Bompfünewerer et al., 2008; Hofacker et al., 1994; Lorenz et al., 2016, 2011; Mückstein et al., 2006). RNAfold was run with default parameters. For RNAplfold, sub-sequences were generated from the input sequences to calculate opening energies (using the parameters -W 210 -u 210). For RNAup, we examined the stochastic interactions between the region 1:30 of each mRNA and 54 non-coding RNAs (using the parameters -b -o). RNAup reports the total interaction between two RNAs as the sum of energy required to open accessible sites in the interacting molecules *G*Δ _*u*_ and the energy gained by subsequent hybridisation *G*Δ _*h*_ (Mückstein et al., 2006). For the interactions between each mRNA and 54 non-coding RNAs, we chose the most stable mRNA:ncRNA pair to report an inappropriate mRNA:ncRNA interaction, i.e. the pair with the strongest hybridisation energy, (Δ*G*_*h*_.) _*min*_.

CAI, tAI and CC were calculated using the reference weights from Sharp and Li (Sharp and Li, 1987), Tuller et al. (Tuller et al., 2010) and Ang et al. (Ang et al., 2016), respectively. Translation elongation rate was predicted using Iχnos(Tunney et al., 2018) trained with ribosome profiling data (SRR7759806 and SRR7759807) (Mohammad et al., 2019).

### Coarse-grained simulation

Our experiments showed a diminishing trend on protein production beyond a certain opening energy (Fig 4). To explain this, we performed a coarse grained simulation using constructs with increasing opening energy on a simulated cellular system. Despite being less precise than fine grained methods such as *ab initio* and molecular dynamics, coarse grained simulations often give similar results, with an added advantage of being scalable to very large systems.

To set the simulation, we binned the opening energies between 2 and 32 in intervals of two, with each bin representing a ‘reporter plasmid construct’ whose opening energy is the mean of the bin. For each construct, the ‘technical replicates’ were generated by allowing slight variations on the mean opening energy of the bin. This is to model variation between replicates, and the discrepancies between the estimated and the actual opening energies *in vivo*. For each round of transcription, mRNA copies were randomly generated from 30 to 60 plasmid DNA copies (Gomes et al., 2020; Held et al., 2003; Rosano and Ceccarelli, 2014). We chose an optimum opening energy of 12 kcal/mol or less for translation. However, this is probabilistic which occasionally allowed protein production from higher opening energy transcripts. We allowed mRNA to decay probabilistically when a mRNA molecule is translated for more than 10 rounds.

We also set a threshold of protein tolerance to be 1,000,000 copies where the copy numbers of endogenous proteins are usually less than 10,000 (Taniguchi et al., 2010), beyond which there is a sporadic death of cells. However, in this simulation, the chances of staying viable and reproducing are higher than death, and cells grow steadily. This threshold also simulated random but low cell deaths in the experiment, without setting an extra variable.

To limit the computational complexity, our coarse-grained simulations used lower constants and iterations. Initialising with 100 cells, the algorithm was set to terminate either after 10,000 iterations or when the total number of cells becomes zero. After termination, the total number of proteins and cells for each construct were taken from the endpoints. To imitate ‘biological replicates’, we repeated the above simulation three times with different random numbers, which provides slightly different initial conditions for each experiment.

### TIsigner development

Finding a synonymous sequence with a maximum accessibility is a combinatorial problem that spans a vast search space. For example, for a protein-coding sequence of nine codons, assuming an average of 3 synonymous codons per amino acid, we can expect a total of 19,682 unique synonymous coding sequences. This number increases rapidly with increasing numbers of codons. Heuristic optimisation approaches are preferred in such situations because the search space can be explored more efficiently to obtain nearly optimal solutions.

To optimise the accessibility of a given sequence, TIsigner uses a simulated annealing algorithm (Brownlee, 2011; Ingber, 2000; Keith et al., 2002; Kirkpatrick et al., 1983), a heuristic optimisation technique based on the thermodynamics of a system settling into a low energy state after cooling. Simulated annealing algorithms have been used to solve many combinatorial optimisation problems in bioinformatics. For example, we previously applied this algorithm to align and predict non-coding RNAs from multiple sequences (Lindgreen et al., 2007). Other studies use this algorithm to find consensus sequences (Keith et al., 2002), optimise ribosome binding sites (Salis et al., 2009) and predict mRNA foldings (Gaspar et al., 2013) using minimum free energy models.

According to statistical mechanics, the probability *p*_*i*_ of a system occupying energy state,*E*_*i*_ with, temperature *T*, follows a Boltzmann distribution of the form 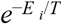, which gives a set of probability mass functions along every point *i* in the solution space. Using a Markov chain sampling, these probabilities are sampled such that each point has a lower temperature than the previous one. As the system is cooled from high to low temperatures (*T*→0), the samples converge to a minimum of *E*, which in many cases will be the global minimum (Keith et al., 2002). A frequently used Markov chain sampling technique is Metropolis-Hastings algorithm in which a ‘bad’ *E*_2_ move from initial state *E*_1_ such that *E*_2_ >*E* _1_, is accept*R*e1d≥if (0,) *p*_2_/ *p*_1_, where (0,)*R*i1s a uniformly random number between 0 and 1.

In our implementation, each iteration consists of a move that may involve multiple synonymous codon substitutions. The algorithm begins at a high temperature where the first move is drastic, synonymous substitutions occur in all replaceable codons. At the end of the first iteration, a new sequence is accepted if the opening energy is smaller than that of the input sequence. However, if the opening energy of a new sequence is greater than that of the input sequence, acceptance depends on the Metropolis-Hastings criteria. The accepted sequence is used for the next iteration, which repeats the above process. As the temperature cools, the moves get milder with fewer synonymous codon changes (Supplementary Fig S6A). Simulated annealing stops upon reaching a near-optimum solution.

For the web version of TIsigner, the default number of replaceable codons is restricted to the first nine codons. However, this default setting can be reset to range from the first four to nine codons, or the full length of the coding sequence. Since the accessibility of a fixed region is optimised, this process only takes O(1) time (Supplementary Fig S7). Furthermore, TIsigner runs multiple simulated annealing instances, in parallel, to obtain multiple possible sequence solutions.

When users select *T7lac* promoter as the 5′UTR, they can adjust ‘Expression Score’, that is calculated based on the PSI:Biology dataset (see below). This allows them to tune the expression level of a target gene. In contrast, when users input a custom 5′UTR sequence, they only have the option to either maximise or minimise expression.

To implement ‘Expression Score’, the posterior probabilities of success for input and optimised sequences are evaluated using the following equations from Bayesian statistics:

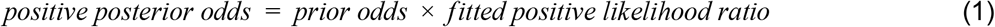

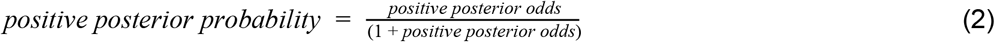

The fitted positive likelihood ratios in equation (1) were obtained from the following 4-parametric logistic regression equation:

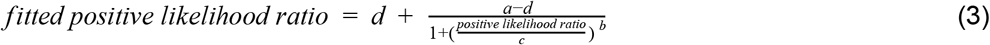

with parameters a, b, c, and d. The prior probability was set to 0.49, which is the proportion of ‘Expressed’ (N=21,046) divided by ‘Cloned’ (N=42,774) of the PSI:Biology targets reported as of 28 June 2017 (http://targetdb.rcsb.org/metrics/). Posterior probabilities were scaled as percentages to score the input and optimised sequences.

The presence of terminator-like elements (Chen et al., 2013) in the protein-coding region may result in expression of truncated mRNAs due to early transcription termination. Therefore, we implemented an optional check for putative terminators in the input and optimised sequences by cmsearch (INFERNAL version 1.1.2) (Nawrocki and Eddy, 2013) using the covariance models of terminators from RMfam (Gardner and Eldai, 2015; Kalvari et al., 2018). We also allow users to filter the output sequences for the presence of restriction sites. Restriction modification sites (AarI, BsaI, and BsmBI) are avoided by default.

Besides *E. coli*, users can choose *S. cerevisiae, M. musculus* or ‘Other’ as the expression host. The regions for optimising accessibility are −7:89, −8:11 and −24:89 for *S. cerevisiae, M. musculus* and ‘Other’, respectively (Fig 1 and Supplementary Fig S1). When users choose ‘Custom’ for expression host, the region for optimising accessibility becomes customisable.

### Sequence optimisation

We submitted the PSI:Biology targets that failed to be expressed (N=2,650) to the ExpOptimizer web server from NovoPro Bioscience (https://www.novoprolabs.com/tools/codon-optimization). A total of 2,573 sequences were optimised. The target sequences were also optimised using a local version of COOL (Chung and Lee, 2012) and TIsigner using default settings. We also ran a random synonymous codon substitution as a control for these 2,573 sequences.

### GFP assay

Plasmids were constructed using the MIDAS Golden Gate cloning system (Supplementary Methods) (van Dolleweerd et al., 2018). BL21(DE3)pLysS competent *E. coli* cells (Invitrogen) were transformed with plasmids and grown overnight on Luria-Bertani (LB) agar plates containing spectinomycin (50 µg/ml) and chloramphenicol (25 µg/ml). Single colonies were picked and inoculated into 3 ml LB broth containing the same antibiotics, and cultures were grown for 18 hours at 37°C, 200 rpm. Cultures were diluted with fresh media at 1:20 and grown at 37°C, 200 rpm, until reaching the mid-logarithmic growth phase (optical densities at 600 nm (OD_600_) of ∼0.3). Of each culture, 20 µl was seeded into 96-well plates containing 180 µl LB broth supplemented with antibiotics and isopropyl-β-D thiogalactopyranoside (IPTG) (1 mM final concentration) per well. Fluorescence intensities and ODs were measured in a black, flat, clear bottom 96-well plate with lid (CELLSTAR, Greiner) using a FLUOstar Omega plate reader (BMG Labtech) equipped with an excitation filter (band pass 485-12) and an emission filter (band pass 520) for GFP and excitation filter (band pass 484) and an emission filter (band pass 610-10) for mScarlet-I. The plate was incubated at 37°C with “meander corner well shaking” at 300 rpm for 7 hours measuring fluorescence and ODs every 10 minutes. Fluorescence was measured in a 2 mm circle recording the average of 8 measurements per well. Average values of technical replicates were calculated and normalised to the mScarlet-I second reporter, and then to the normalised value of the GFP variant with the highest opening energy (21.68 kcal/mol). Normalised fluorescence values were obtained from the average values of biological replicates (Supplementary Table S2).

### Luciferase assay

BL21Star(DE3) competent cells (Invitrogen) were transformed with plasmids and grown overnight at 37°C on LB agar plates containing 50 µg/ml spectinomycin. Single colonies were picked and inoculated into 5 ml LB broth (50 µg/ml spectinomycin) and grown for 18 hours at 37°C, 200 rpm. Bacterial cultures were diluted with fresh media at 1:20 and grown at 37°C, 200 rpm, up to a mid-logarithmic phase (OD_600_ of ∼0.4). The cultures were split and induced with IPTG at a final concentration of 0.25 mM (or uninduced as controls), and seeded into a white, flat, clear bottom 96-well white plate with lid (Costar, Corning), 150 µl per well, in triplicates. Cells were incubated in a FLUOstar Omega Microplate Reader (BMG LABTECH) for 90 minutes at 25°C, 200 rpm, and OD_600_ was measured every 15 minutes (over 7 cycles). Cells were harvested by centrifugation at 3000 ×g, for 10 minutes, at 20°C. Supernatants were removed. As the substrate can penetrate into cells, 50 µl of coelenterazine h (Promega) was added to living cells to minimise sample processing steps and variability (Fuhrmann et al., 2004; Lorenz et al., 1996). Luminescence was measured (λ_em_ = 475 nm) in a Clariostar microplate reader (BMG LABTECH) at 25°C every 2 minutes (over 11 cycles). Average values of technical replicates were calculated and normalised to the wild-type. Normalised luminescence values were obtained from the average values of biological replicates (Supplementary Table S2).

### Statistical analysis

AUC and Gini importance scores were calculated using scikit-learn (version 0.20.2) (Pedregosa et al., 2011). The 95% confidence intervals for AUC scores were calculated using DeLong’s method (DeLong et al., 1988). Spearman’s correlation coefficients and Kolmogorov-Smirnov statistics were calculated using Pandas (version 0.23.4) (McKinney, 2010) and scipy (version 1.2.1) (Millman and Aivazis, 2011; Oliphant, 2007), respectively. Positive likelihood ratios with 95% confidence intervals were calculated using the bootLR package (Marill et al., 2017; R Core Team, 2019). The P-values of multiple testing were adjusted using Bonferroni’s correction and reported to machine precision. Plots were generated using Matplotlib (version 3.0.2) (“Matplotlib: A 2D Graphics Environment - IEEE Journals & Magazine,” n.d.) and Seaborn (version 0.9.0) (Waskom et al., 2018).

## Code and data availability

Our code and data can be found in our GitHub repository (https://github.com/Gardner-BinfLab/TIsigner_paper_2019). These include the scripts and Jupyter notebooks to reproduce our results and figures. The source code of TIsigner is available at https://github.com/Gardner-BinfLab/TISIGNER-ReactJS. The public web version of this tool runs at https://tisigner.com/tisigner. The experimental data, analysis and results are available at https://github.com/bkb3/TIsignerExperiment/tree/master/Jupyter and an interactive version of results are available at https://bkb3.github.io/TIsignerExperiment/.

## ACKNOWLEDGEMENTS

We thank Professor Ivo Hofacker for fruitful discussions at the Benasque RNA Meeting, and Dr Ronny Lorenz for helpful discussions about RNAplfold. We are grateful to the members of the Biomolecular Interaction Centre at the University of Canterbury for supporting this research. We thank New Zealand eScience Infrastructure for providing high performance computing resources. This work was supported in part by the Ministry of Business, Innovation and Employment [MBIE Smart Idea grant: UOOX1709 and MBIE Data Science Programmes grant: UOAX1932] and the Royal Society of New Zealand Te Apārangi [Marsden grant: 19-UOO-040].

## AUTHOR CONTRIBUTIONS

C.S.L. and P.P.G. conceived the work; C.S.L. contributed RNA accessibility analyses; B.K.B. performed the coarse-grained simulation and developed the TIsigner web server; C.D., D.M.R., and A.C. constructed the plasmids, performed the GFP assay, and the luciferase assay, respectively. C.S.L. and B.K.B. analysed the data and drafted the manuscript. All authors reviewed, edited and approved the manuscript.

