## Supplementary Fig, Supplementary Methods for "Protein yield is tunable by synonymous codon changes of translation initiation sites"

### SUPPLEMENTARY FIGURES

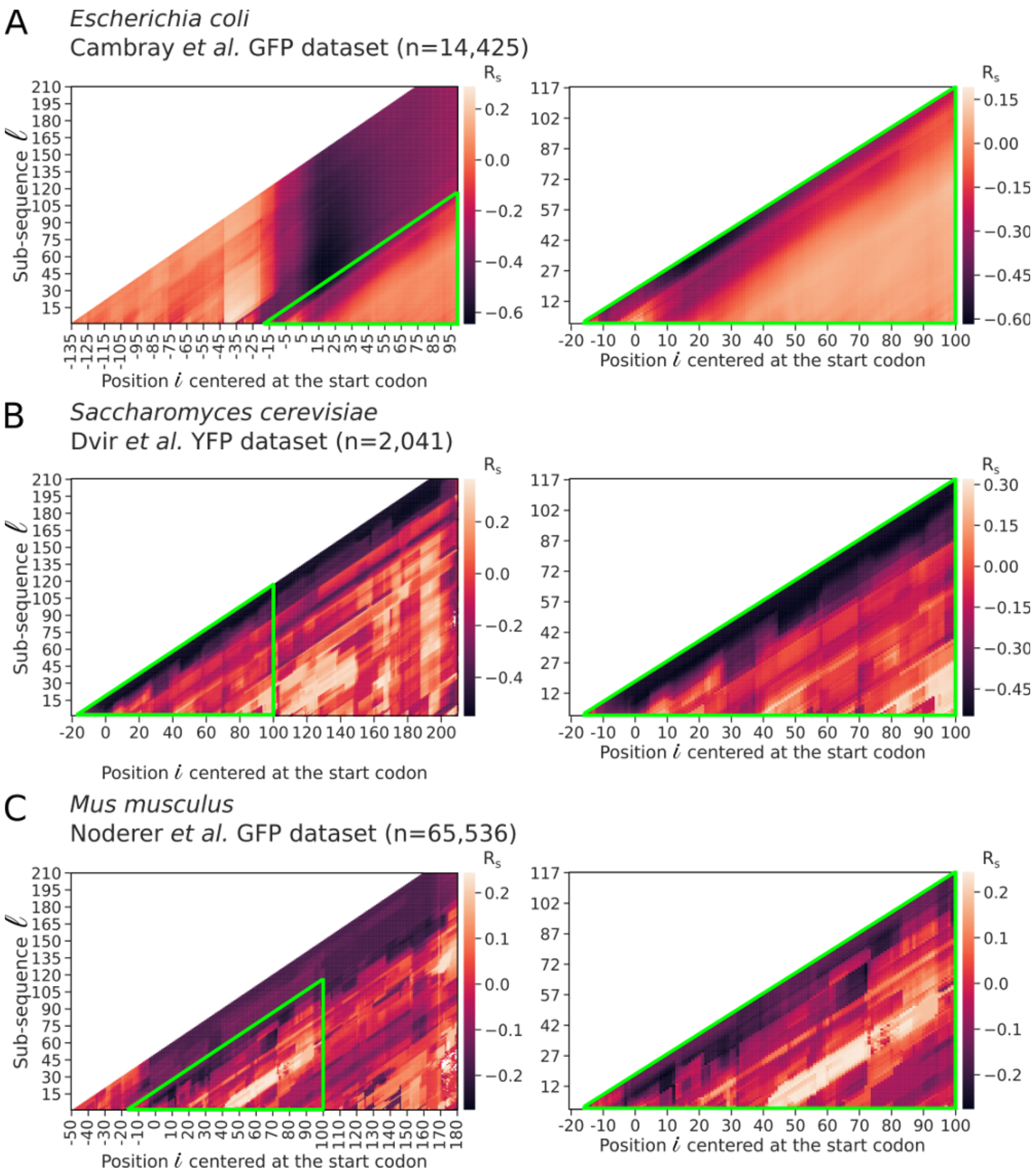


**Fig S1. Heatmaps of correlations between opening energy and protein abundance for each of the sub-sequence regions (related to Fig 1).** Green unfilled triangles indicate the regions before and after scaling (left and right panels, respectively). **(A)** For *E. coli*, we used a representative GFP expression dataset from Cambray et al. (2018) (Cambray et al., 2018). The reporter library consists of GFP fused in-frame with a library of 96-nt upstream sequences (n=14,425). **(B)** For *S. cerevisiae*, we used a YFP expression dataset from Dvir et al. (2013) (Dvir et al., 2013). The YFP reporter library consists of 2,041 random decameric nucleotides inserted at the upstream of YFP start codon. **(C)** For *M. musculus*, we used the GFP expression dataset from Noderer et al. (2014) (Noderer et al., 2014). The GFP reporter library consists of 65,536 random hexameric and dimeric nucleotides inserted at the upstream and downstream of GFP start codon, respectively. $R_s$, Spearman's rho.

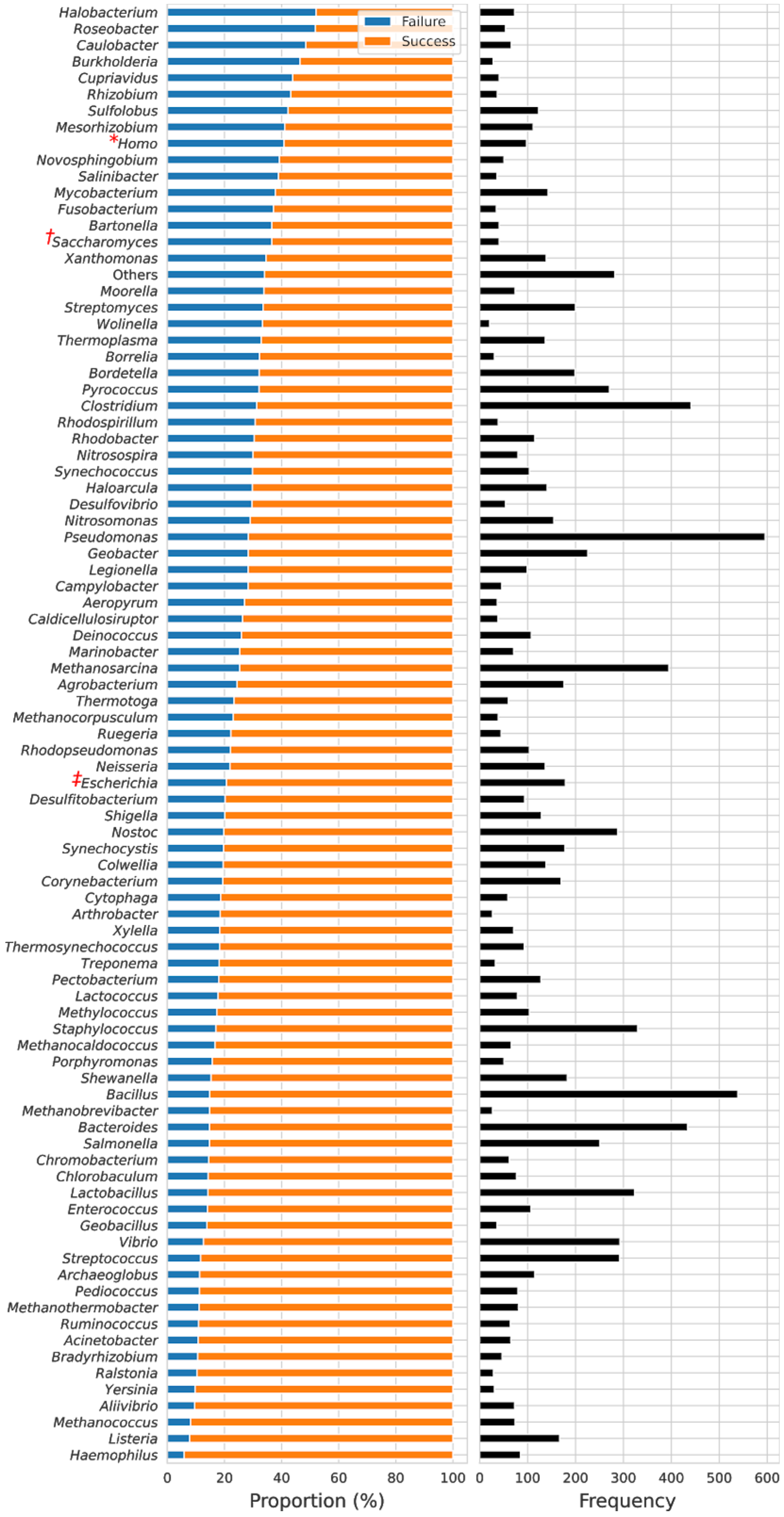


**Fig S2. Expression outcomes of the PSI:Biology targets in *E. coli*.** A total of 11,430 PSI:Biology targets from over 189 species were analysed in this study (n=8,780 and 2,650, 'success' and 'failure' groups, respectively) (Acton et al., 2005; Chen et al., 2004; Seiler et al., 2014). Genera with at least 20 target genes are shown and the remaining as 'Others'. The top three PSI:Biology targets are from four *Pseudomonas*, five *Bacillus* and six *Clostridium* species. Red asterisk, obelisk and diesis indicate *Homo sapiens*, *S. cerevisiae* and *E. coli*, respectively. These target genes were inserted into the pET21_NESG expression vector, in which the promoter and fusion tag are *T7lac* and C-terminal His tag, respectively.

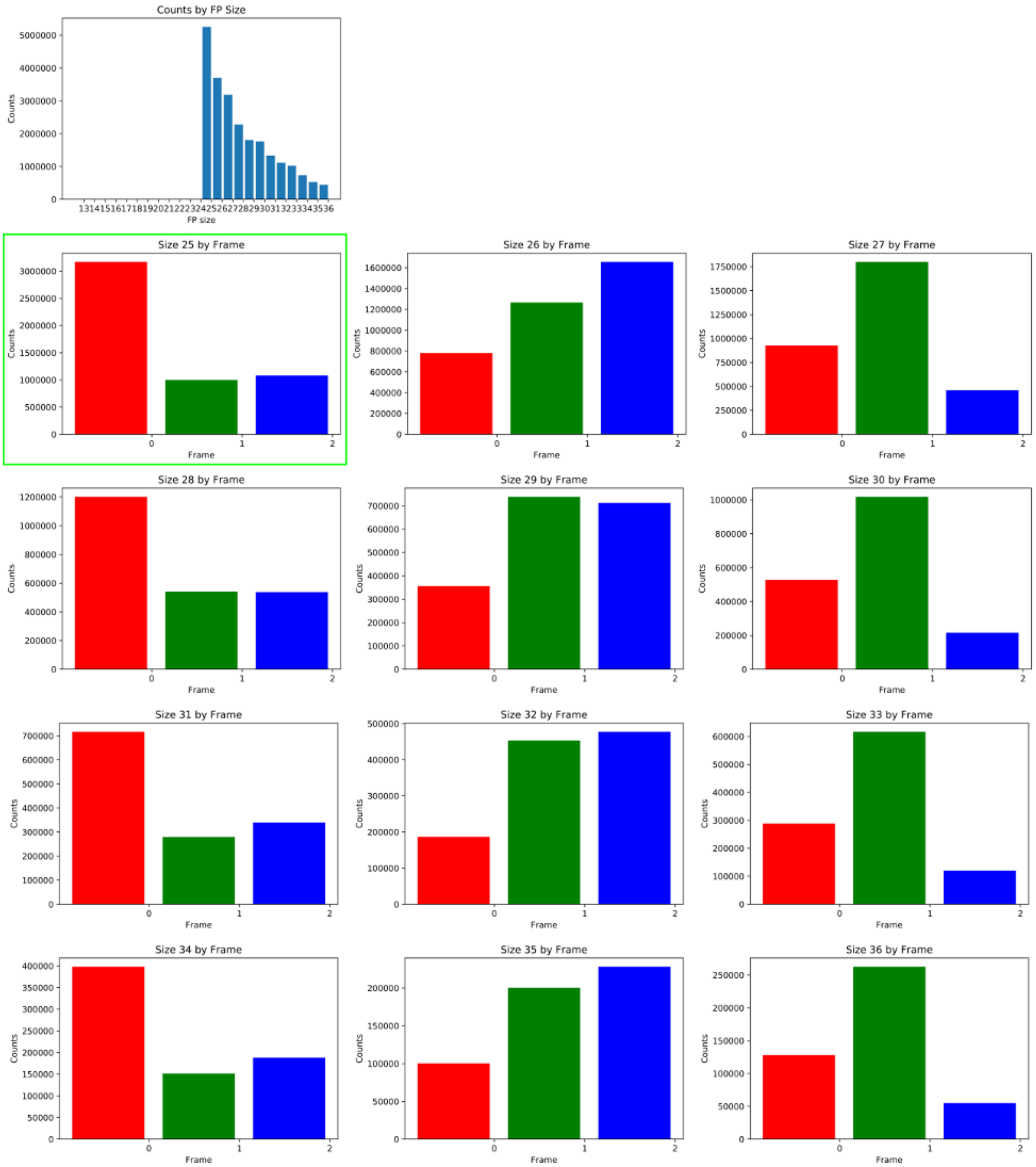


**Fig S3. Distributions of ribosome footprints by fragment sizes and reading frames.** Ribosome profiling data [SRR7759806 and SRR7759807 (Mohammad et al., 2019)] were aligned to *S. cerevisiae* transcriptome (see https://github.com/Gardner-BinfLab/TIsigner_paper_2019). SAM alignment files were merged, and ribosome footprints which were mapped to each frame were enumerated. Ribosome footprints of 25 nt in length have the greatest proportion of footprints mapped to frame zero (green unfilled rectangle). These 25-nt footprints were used to train a neural network model (Tunney et al., 2018) in order to predict the translation elongation rates of the PSI:Biology targets. FP, footprints.

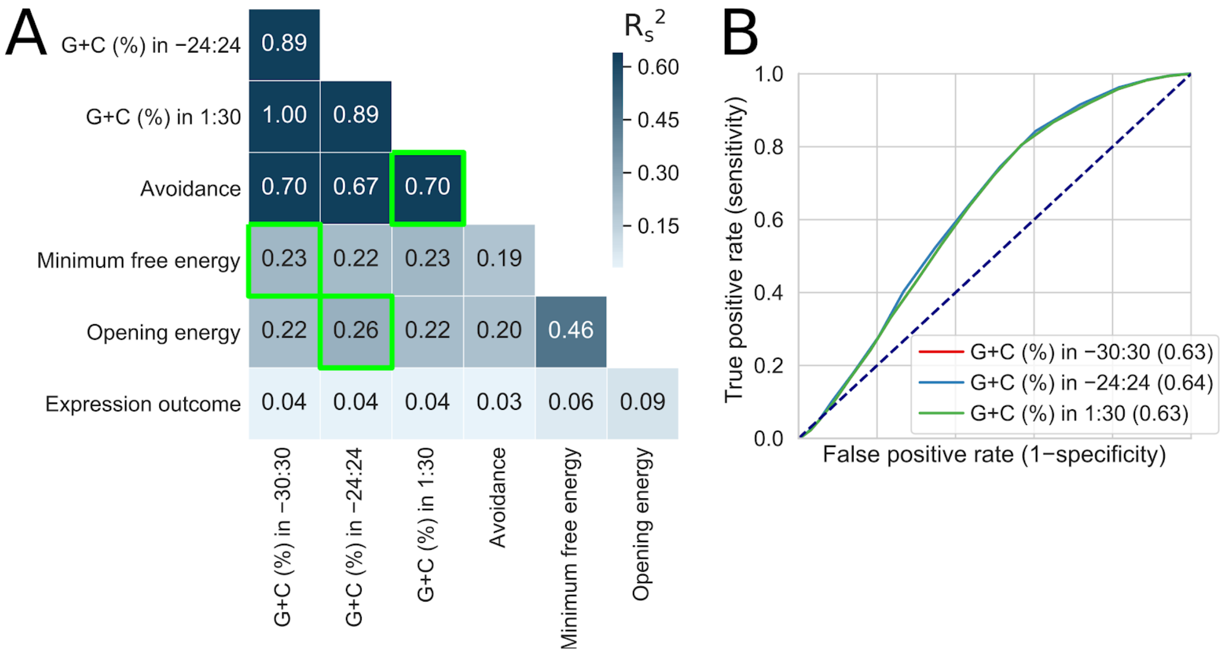


**Fig S4. Analysis of the local G+C contents (related to Fig 1). (A)** The G+C contents in the regions −24:24 and −30:30 weakly correlate with opening energy and minimum free energy, respectively. Green unfilled squares indicate squared Spearman's correlations ($R_s^2$) between the local G+C contents and the corresponding local features. **(B)** The local G+C contents show a similar prediction accuracy (AUC scores shown in parentheses).

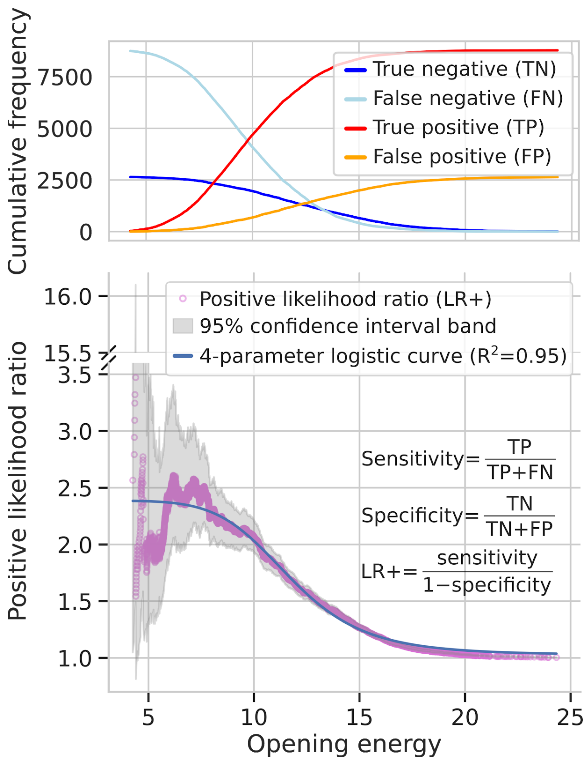


**Fig S5. Opening energy of 10 or below at the region −24:24 is about two times more likely to come from the target genes that are successfully expressed than those that failed (with 95% confidence).** Cumulative frequency distributions of the true positive and false positive (less than type), and true negative and false negative (more than type) derived from the ROC analysis in Fig 2A (left panel, opening energy −24:24). These values were used to estimate positive likelihood ratios with 95% confidence intervals using 10,000 bootstrap replicates. The estimated ratios and/or confidence intervals are inaccurate at low numbers of true positives or true negatives. Therefore, a four-parameter logistic curve was fitted to the positive likelihood ratios. Fitted values are useful to estimate the posterior probability of protein expression.

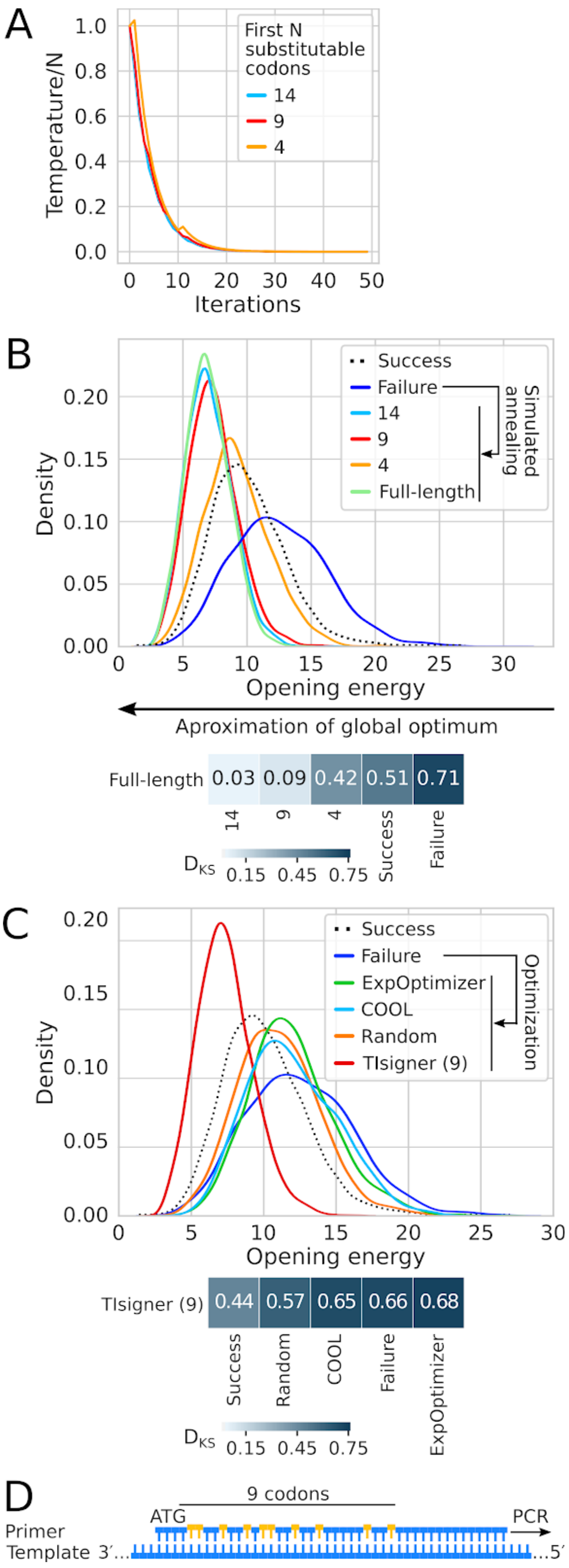


**Fig S6. Accessibility of translation initiation sites can be increased by synonymous codon substitution within the first nine codons using simulated annealing. (A)** Schedules in simulated annealing. The ratio of temperature to the number of the first N substitutable codons decreases exponentially with increasing number of iterations. **(B)** Accessibility of translation initiation sites increases with increasing number of the first N replaceable codons. The PSI:Biology targets that failed to be expressed were optimised using simulated annealing (n=2,650). The Kolmogorov-Smirnov distance between the distributions of '9' and 'full-length' was significantly different but sufficiently close ($D_{KS}$=0.09, $P<10^{-7}$), indicating that optimisation of the first nine codons can achieve nearly optimum accessibility. For comparison, the distribution of the PSI:Biology targets that were successfully expressed are shown (n=8,780). See also Table S1. **(C)** Accessibility of translation initiation sites can be increased indirectly using the existing gene optimisation tools and random synonymous codon substitution. 'TIsigner (9)' refers to the default settings of our tool, which allows synonymous substitutions up to the first nine codons (as above). See also Table S1. **(D)** Accessibility of translation initiation sites can be optimised using PCR cloning. The forward primer should be designed according to TIsiger optimised sequences. For example, using a nested PCR approach, the optimised sequence can be produced using the forward primer designed with appropriate mismatches (gold bulges) to amplify the amplicon from the initial PCR reaction.

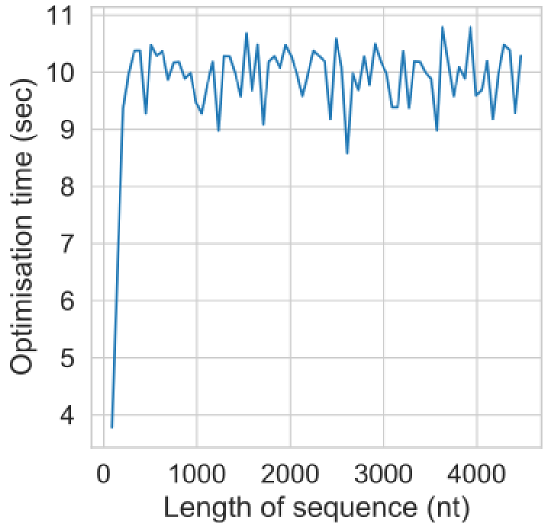


**Fig S7. Sequence length does not affect software performance because only a fixed region is taken into account during optimisation (O(1) time).**

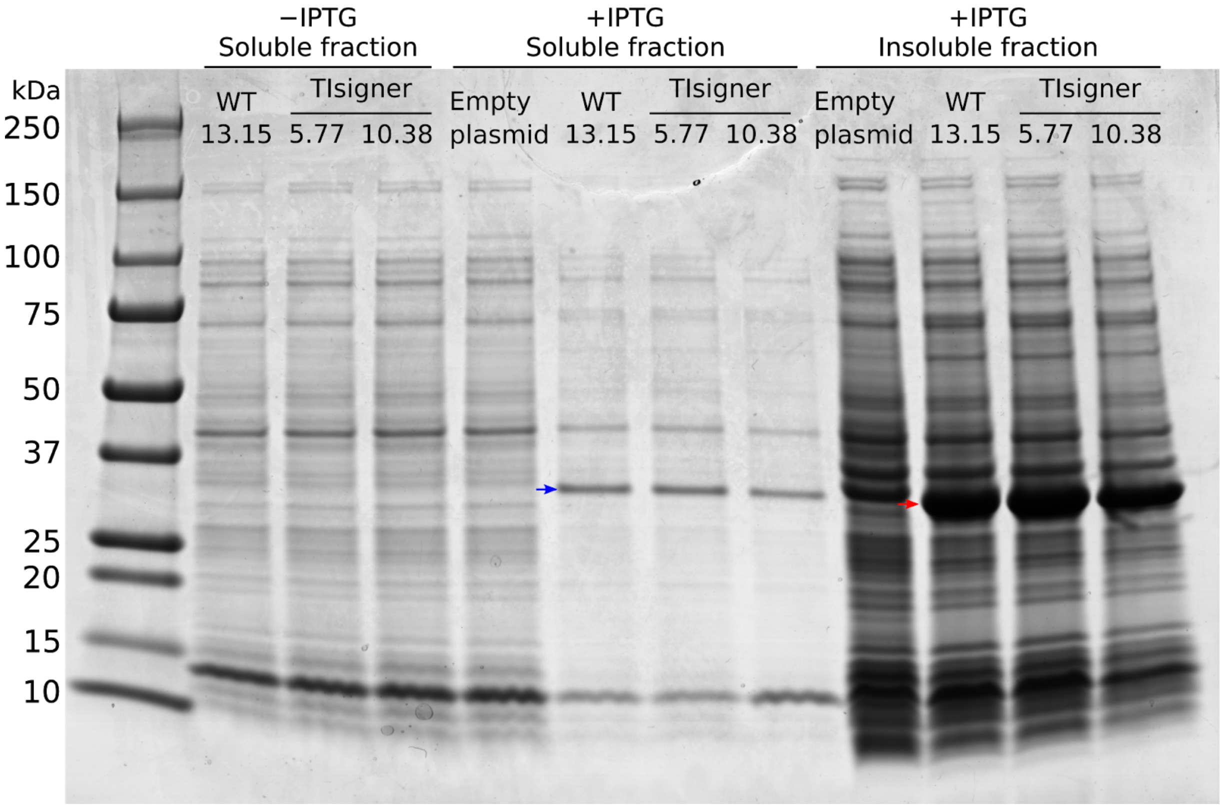


**Fig S8. SDS-PAGE gel shows the protein bands of *Renilla* luciferase (RLuc) in the soluble and insoluble fractions of BL21Star(DE3) lysates.** Selected bacterial clones were grown at 25 °C, 200 RPM. The solubilities of wildtype (WT) RLuc and designed variants were compared after 4-hour IPTG induction. The blue and red arrows (~36kDa) indicate that RLuc was poorly soluble. No RLuc protein bands were detected from the uninduced cultures and IPTG-induced negative control (empty vector control that lacks Rluc gene and *T7lac* promoter).

### SUPPLEMENTARY METHODS

#### Cloning of TIsigner variants of GFP and Luciferase

The cloning of TIsigner sequence variants for *E. coli* expression was performed using the MIDAS Golden Gate cloning system (van Dolleweerd et al., 2018). As with other Golden Gate assembly (GGA) systems, MIDAS is a modular, hierarchical DNA assembly system that uses the Type IIS restriction enzymes AarI, BsaI and BsmBI to assemble genes, transcription units and other devices from basic parts, and subsequently enables multiple devices to be assembled together on a single plasmid.

As per MIDAS, basic parts such as promoters, coding sequences and terminators were amplified by PCR or ordered as synthetic polynucleotide sequences from gene synthesis companies. The basic parts are listed in Table S3. Protocols for the GGA reactions are as described in van Dolleweerd et al., 2018 (van Dolleweerd et al., 2018).

*N-terminal region of GFP (GFPN)*

Overlapping oligonucleotide primer pairs corresponding to the first ten codons of each of the *gfp* sequence variants produced by the TIsigner algorithm were ordered from Integrated DNA Technologies (IDT) (see Table S4). Each overlapping pair of primers was annealed together and used as a template for amplification by Q5 polymerase (New England Biolabs) to create a double-stranded DNA product spanning the N-terminal region of GFP (GFPN; codons 1 to 10; see Fig S9), and with MIDAS [CCAT] prefix and [GTTG] suffix nucleotides.

*C-terminal region of GFP (GFPC)*

The C-terminal region of GFP (designated GFPC), spanning codons 11 to 238 of the native *gfp* of *Aequoria victoria*, was synthesized (GeneArt) with flanking BsmBI recognition sites, and with the [GTTG] MIDAS prefix (compatible with the [GTTG] suffix on the GFPN part) and [GCTT] suffix nucleotides.

*N-terminal region of luciferase (RLucN)*

Overlapping oligonucleotide primer pairs corresponding to the first ten codons of each of the luciferase sequence variants generated by the TIsigner algorithm were ordered from Integrated DNA Technologies (IDT) (see Table S5). Each overlapping pair of primers was annealed together and used as a template for amplification by Q5 polymerase (New England Biolabs) to create a double-stranded DNA product spanning the N-terminal region of luciferase (RLucN; codons 1 to 10), and with a MIDAS [CCAT] prefix and an [AGGA] suffix.

*C-terminal region of luciferase (RLucC)*

The C-terminal region of luciferase (designated RLucC), spanning codons 11 to 311 of the native luciferase of *Renilla reniformis*, was amplified from the full-length, native luciferase sequence (synthesized by GeneArt) using primers that add flanking BsmBI recognition sites, and a MIDAS [AGGA] prefix (compatible with the [AGGA] suffix on the RLucN part) and a [GCTT] suffix.

<u>MIDAS Level-1 cloning of parts</u>

PCR products, purified using commercially available column-based protocols (Macherey-Nagel), or parts produced by gene synthesis were cloned into the MIDAS pML1 vector by BsmBI-mediated Golden Gate assembly (BsmBI-GGA). As per the MIDAS design, BsmBI-GGA into the pML1 vector results in elimination of the BsmBI recognition sites and each part becomes flanked by BsaI recognition sites that cleave at the MIDAS prefix and suffix nucleotides.

In the case of the *Aequoria victoria gfp* TIsigner variants, each GFPN part cloned into the pML1 vector becomes flanked by BsaI recognition sites that are cleaved at the [CCAT] prefix and [GTTG] suffix (Fig S10). Cloning of the GFPC part into the pML1 vector results in a GFPC module flanked by BsaI recognition sites that are cleaved at the [GTTG] prefix and at the [GCTT] suffix (Fig S11).

In the case of the *Renilla reniformis* luciferase TIsigner variants, each RLucN part cloned into the pML1 vector becomes flanked by BsaI recognition sites that are cleaved at the [CCAT] prefix and [AGGA] suffix, while the C-terminal fragment, RLucC, becomes flanked by BsaI recognition sites that generate an [AGGA] prefix and a [GCTT] suffix upon cleavage.

All parts cloned into the pML1 vector were verified by sequencing.

<u>MIDAS Level-2 assembly of devices</u>

Devices were assembled from the cloned Level-1 modules described above, using BsaI-GGA, into the appropriate pML2 vector. As per the MIDAS design, multiple parts can be assembled together, with the position of each part in the assembled device dictated by the compatibility of the prefix and suffix nucleotides flanking each module:

- A *lacI* device was assembled in pML2(+)WR from the single *lacI* genetic element module.
- An *mScarlet-I* device was assembled in pML2(+)BR from *nptII* promoter, *mScarlet-I* CDS and lambda t0 transcription terminator modules.
- Full-length *gfp* devices for each TIsigner variant were assembled in pML2(+)WF from the following modules: *T7lac* promoter, GFPN, GFPC and T7 Tφ transcription terminator. Since the prefix of the GFPC module, [GTTG] (see Fig S11), is identical to the suffix of each GFPN module (see Fig S10) this allows the two modules to be genetically fused so that, together with the *T7lac* promoter and T7 Tφ transcription terminator modules, full-length *gfp* devices are assembled for each variant. The *mScarlet-I* and *gfp* devices were assembled in pML2 vectors of opposite orientation (using the "Reverse" vector pML2(+)BR for *mScarlet-I*, and the "Forward" vector pML2(+)WF for each *gfp* device), so that they will be divergently transcribed once assembled into the expression vector (Level-3, see below).
- In a similar fashion, full-length luciferase devices for each TIsigner variant were assembled in pML2(+)BF from *T7lac* promoter, RLucN, RLucC and T7 Tφ transcription terminator modules.

All cloned devices were verified by restriction mapping and sequencing.

<u>MIDAS Level-3 assembly (construction of the expression plasmids)</u>

*E. coli gfp* expression plasmids were constructed by sequentially loading the *lacI*, *mScarlet-I* and *gfp* devices, using alternating AarI- and BsmBI-GGA reactions, into the MIDAS Level-3 destination plasmid pML3.2, which has the medium copy replication origin from the pET series of vectors in place of the high copy pMB1 replicon of the pML3 destination vector originally

described in van Dolleweerd et al, 2018 (van Dolleweerd et al., 2018). A representative map of an *E. coli* expression plasmid containing all three devices is shown in Fig S12.
For luciferase expression, the intermediate plasmid containing the *lacI* device (described above) was used for assembly of each of the luciferase devices (i.e., no *mScarlet-I* device was added), and a representative map of an *E. coli* plasmid for luciferase expression is shown in Fig S13. The *lacI* and luciferase devices are divergently transcribed from the expression vector.

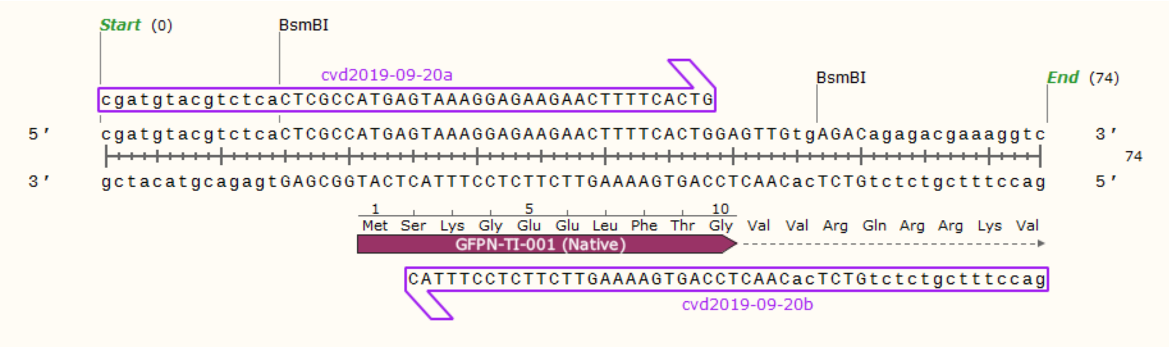


**Fig S9. Strategy for producing a double stranded DNA corresponding to the first ten codons of each of the TIsigner variants of GFP.** The strategy employs a pair of primers that overlap at their 3' ends that, upon annealing together, can be used as a template for Q5 polymerase to generate a double stranded DNA spanning the N-terminal region of GFP (i.e. GFPN). Shown here is the sequence of variant GFPN-001 generated using the overlapping cvd2019-09-20a forward and cvd2019-09-20b reverse primer pair (see Table S5). The resultant double stranded DNA can then be cloned into the MIDAS pML1 vector by digestion with the Type IIS restriction enzyme BsmBI (recognition site CGTCTC(1/5)). The same primer pair strategy was used for producing TIsigner variants of luciferase, albeit with a different [AGGA] suffix. This map was created with SnapGene.

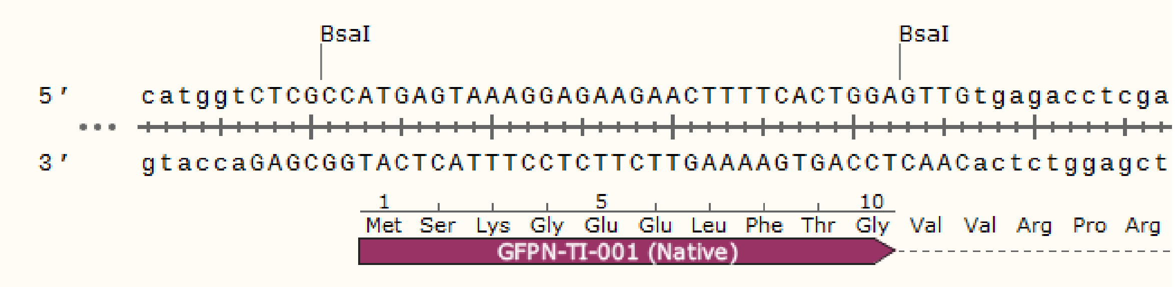


**Fig S10. Structure of GFPN variants cloned into the pML1 vector.** Following BsmBI-mediated cloning into the pML1 vector, each GFPN variant becomes flanked by BsaI recognition sites (GGTCTC(1/5)). BsaI cleaves at the CCAT prefix upstream of the GFPN sequence and at the GTTG suffix (downstream of the GFPN module). For ease of depiction, only the sequences immediately surrounding the cloned GFPN fragment are shown (i.e., not the rest of the pML1 vector). The structure of luciferase RLucN variants is identical, except for having a different [AGGA] suffix sequence.

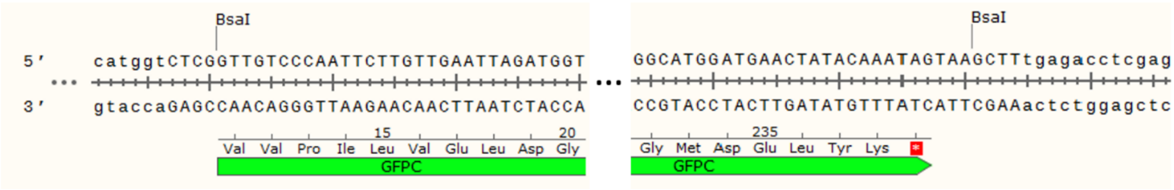


**Fig S11. Structure of the GFPC fragment cloned into the pML1 vector.** Following BsmBI-mediated cloning into the pML1 vector, GFPC becomes flanked by BsaI recognition sites. BsaI cleaves at the [GTTG] prefix (upstream of the GFPC sequence) and at the downstream [GCTT] suffix. For ease of depiction, only sequences around the 5’ and 3’ ends of the GFPC fragment are shown (left- and right-hand sides, respectively). In the case of luciferase, the prefix sequence is [AGGA].

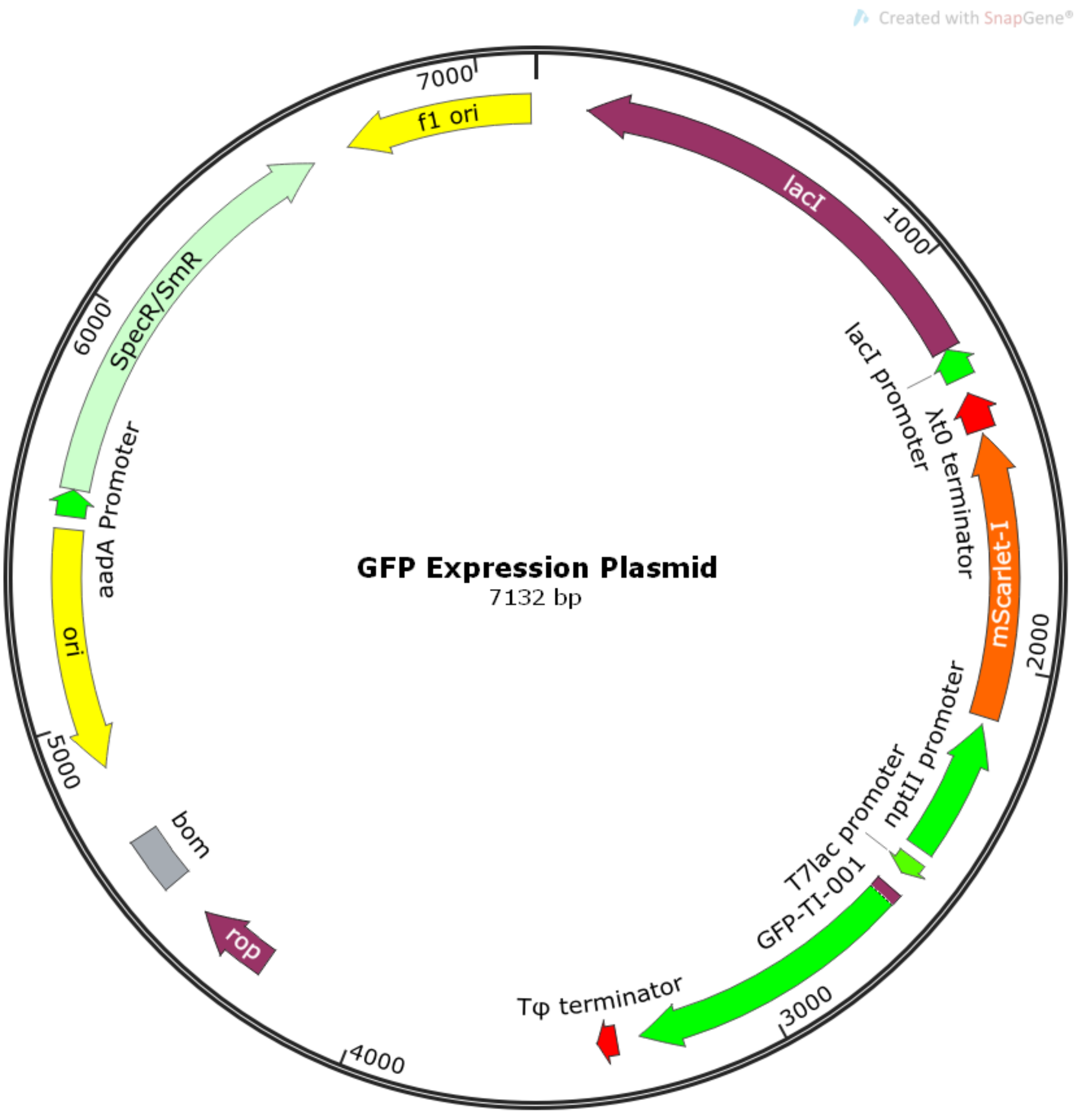


**Fig S12. Structure of GFP expression plasmids.** Map view showing the architecture of MIDAS-assembled plasmids used for expression of GFP TIsigner variants. Expression of each of the gfp variants is controlled by the *T7lac* promoter and oriented such that they are divergently transcribed with respect to the *mScarlet-I* device, which is driven by the *nptII* promoter. The devices for *lacI*, *mScarlet-I* and *gfp* were loaded sequentially into plasmid pML3.2, which has the medium copy replication origin from the pET series of vectors (*ori-bom-rop*) in place of the high copy pMB1 replicon in the pML3 destination vector described in van Dolleweerd et al, 2018 (van Dolleweerd et al., 2018), and a selectable marker conferring resistance to spectinomycin.

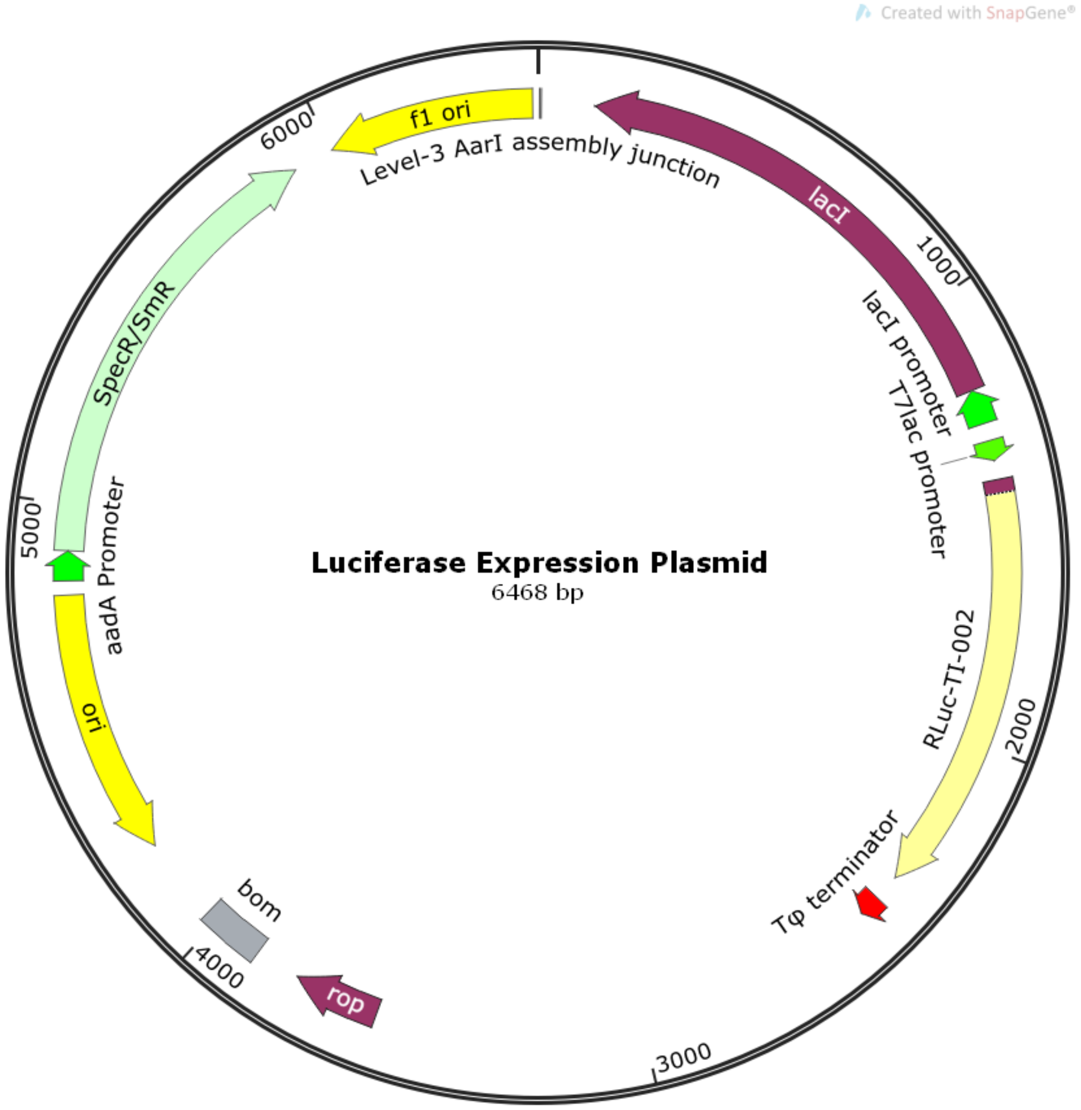


**Fig S13. Structure of luciferase expression plasmids.** Map view showing the architecture of MIDAS-assembled plasmids used for expression of luciferase TIsigner variants. Expression of each of the luciferase variants is controlled by the *T7lac* promoter and oriented such that they are divergently transcribed with respect to the *lacI* device, which is driven by the *lacI* promoter. The devices for *lacI* and luciferase were loaded sequentially into plasmid pML3.2, which has the medium copy replication origin from the pET series of vectors (*ori-bom-rop*) in place of the high copy pMB1 replicon in the pML3 destination vector described in van Dolleweerd et al, 2018 (van Dolleweerd et al., 2018), and a selectable marker conferring resistance to spectinomycin.

**Table S3. MIDAS parts used in this work.** The sequence of each part is shown, with BsmBI recognition sites used for cloning into the MIDAS pML1 vector underlined. MIDAS prefix and suffix nucleotides are highlighted by the bold blue and red text, respectively. The *T7lac* promoter and T7 Tφ transcription terminator (both sequences taken from pET-15b) were ordered as double-stranded DNA gBlocks from IDT. The *lacI* genetic element (with *lacI* promoter in green and lacI coding sequence in blue) was from GeneArt. Other parts were amplified by polymerase chain reaction. For the *mScarlet-I* part, the coding sequence is highlighted by the blue shading. Sequences of all parts were confirmed following cloning into the MIDAS pML1 vector.

| Part | Sequence |
|---|---|
| *T7lac* promoter | cgatgtacgtctcaCTCG**GGAG**CGATCCCGCGAAATTAATACGACTCACTAT<br>AGGGGAATTGTGAGCGGATAACAATTCCCCTCTAGAAATAATTTTGTT<br>TAACTTTAAGAAGGAGATATA**CCAT**tgAGACagagacgaaaggtc |
| *nptII* promoter | cgatgtacgtctcaCTCG**GGAG**ctcgcacgctgccgcaagcactcagggcgcaagggctgct<br>aaaggaagcggaacacgtagaaagccagtccgcagaaacggtgctgaccccggatgaatgtca<br>gctactgggctatctggacaagggaaaacgcaagcgcaaagagaaagcaggtagcttgcagtgg<br>gcttacatggcgatagctagactgggcggtcttatggacagcaagcgaaccggaattgccagctgg<br>ggcgccctctggtaaggttgggaagccctgcaaagtaaactggatggctttcttgccgccaaggatct<br>gatggcgcaggggatcaagatctgatcaagagacaggatactagtggaggaagaaaa**AATG**t<br>gAGACagagacgaaaggtc |
| *lacI* genetic element | cgatgtacgtctcaCTCG**GGAG**CGTCGAGATCCCGGACACCATCGAATGG<br>CGCAAAACCTTTCGCGGTATGGCATGATAGCGCCCGGAAGAGAGTC<br>AATTCAGGGTGGTGAATGTGAAACCAGTAACGTTATACGATGTCGCA<br>GAGTATGCCGGTGTCTCTTATCAGACCGTTTCCCGCGTGGTGAACC<br>AGGCCAGCCACGTTTCTGCGAAAACGCGGGAAAAAGTGGAAGCGG<br>CGATGGCGGAGCTGAATTACATTCCCAACCGCGTGGCACAACAACT<br>GGCGGGCAAACAGTCGTTGCTGATTGGCGTTGCCACCTCCAGTCTG<br>GCCCTGCACGCGCCGTCGCAAATTGTCGCGGCGATTAAATCTCGCG<br>CCGATCAACTGGGTGCCAGCGTGGTGGTGTCGATGGTAGAACGAAG<br>CGGCGTCGAAGCCTGTAAAGCGGCGGTGCACAATCTTCTCGCGCAA<br>CGCGTCAGTGGGCTGATCATTAACTATCCGCTGGATGACCAGGATGC<br>CATTGCTGTGGAAGCTGCCTGCACTAATGTTCCGGCGTTATTTCTTG<br>ATGTCTCTGACCAGACACCCATCAACAGTATTATTTTCTCCCATGAAG<br>ACGGTACGCGACTGGGCGTGGAGCATCTGGTCGCATTGGGTCACC<br>AGCAAATCGCGCTGTTAGCGGGCCCATTAAGTTCTGTCTCGGCGCG<br>TCTGCGTCTGGCTGGCTGGCATAAATATCTCACTCGCAATCAAATTCA<br>GCCGATAGCGGAACGGGAAGGCGACTGGAGTGCCATGTCCGGTTT<br>TCAACAAACCATGCAAATGCTGAATGAGGGCATCGTTCCCACTGCGA<br>TGCTGGTTGCCAACGATCAGATGGCGCTGGGCGCAATGCGCGCCAT<br>TACCGAGTCCGGGCTGCGCGTTGGTGCGGATATCTCGGTAGTGGGA<br>TACGACGATACCGAAGACAGCTCATGTTATATCCCGCCGTTAACCAC<br>CATCAAACAGGATTTTCGCCTGCTGGGGCAAACCAGCGTGGACCGC<br>TTGCTGCAACTCTCTCAGGGCCAGGCGGTGAAGGGCAATCAGCTGT<br>TGCCCGTGTCACTGGTGAAAAGAAAAACCACCCTGGCGCCCAATAC<br>GCAAACCGCCTCTCCCCGCGCGTTGGCCGATTCATTAATGCAGCTG<br>GCACGACAGGTTTCCCGACTGGAAAGCGGGCAGTGAGCGCAACGC<br>AATTAATGTAAGTTAGCTCACTCATTAGGCACCGGGATCTCGACCGAT |

| | |
|---|---|
| | GCCCTTGAGAGCCTTCAACCCAGTCAGCTCCTTCCGGTGGGCGCG<br>GGGCATGACTA**CGCT**tgAGACagagacgaaaggtc |
| *mScarlet-I* coding sequence (CDS) | cgatgtacgtctcaCTCG**AATG**gtgagcaagggcgaggcagtgatcaaggagttcatgcggtt<br>caaggtgcacatggagggctccatgaacggccacgagttcgagatcgagggcgagggcgaggg<br>ccgcccctacgagggcacccagaccgccaagctgaaggtgaccaagggtggccccctgcccttct<br>cctgggacatcctgtcccctcagttcatgtacggctccagggccttcatcaagcaccccgccgacatc<br>cccgactactataagcagtccttccccgagggcttcaagtgggagcgcgtgatgaacttcgaggac<br>ggcggcgccgtgaccgtgacccaggacacctccctggaggacggcaccctgatctacaaggtga<br>agctccgcggcaccaacttccctcctgacggccccgtaatgcagaagaagacaatgggctgggaa<br>gcgtccaccgagcggttgtaccccgaggacggcgtgctgaagggcgacattaagatggccctgcg<br>cctgaaggacggcggccgctacctggcggacttcaagaccacctacaaggccaagaagcccgtg<br>cagatgcccggcgcctacaacgtcgaccgcaagttggacatcacctcccacaacgaggactaca<br>ccgtggtggaacagtacgaacgctccgagggccgccactccaccggcggcatggacgagctgta<br>caagtaa**GCTT**tgAGACagagacgaaaggtc |
| Phage T7 Tφ transcription terminator | cgatgtacgtctcaCTCG**GCTT**CAAAGCCCGAAAGGAAGCTGAGTTGGCT<br>GCTGCCACCGCTGAGCAATAACTAGCATAACCCCTTGGGGCCTCTAA<br>ACGGGTCTTGAGGGGTTTTTTGCTGAAAGGAGGAACTATATCCGGAT<br>**CGCT**tgAGACagagacgaaaggtc |
| Lambda t0 transcription terminator | cgatgtacgtctcaCTCG**GCTT**ggactcctgttgatagatccagtaatgacctcagaactccatct<br>ggatttgttcagaacgctcggttgccgccgggcgtttttattggtgagaatccaagctagcttgg**CGC**<br>**T**tgAGACagagacgaaaggtc |

**Table S4. Oligonucleotide primer pairs for constructing TIsigner variants of *gfp*.** The sequences of each forward and reverse primer pair used for constructing each of the *gfp* TIsigner variants is shown. The start codon in each of the forward primers is shaded yellow. BsmBI recognition sites (used for Golden Gate assembly into the MIDAS pML1 vector) are underlined. The MIDAS prefix [CCAT] and suffix [GTTG] (reverse-complement = CAAC) for the GFPN modules are highlighted in bold blue and red, respectively.

| GFPN TIsigner ID | Oligonucleotide Primer Pair | Primer Sequences (5' to 3') |
|---|---|---|
| GFPN-001 | cvd2019-09-20a | cgatgtacgtctcaCTCG**CC**ATGAGTAAAGGAGAAGAACTTTTCACTG |
| | cvd2019-09-20b | gacctttcgtctctGTCTca**CAAC**TCCAGTGAAAAGTTCTTCTCCTTTAC |
| GFPN-002 | cvd2019-09-21a | cgatgtacgtctcaCTCG**CC**ATGTCGAAGGGTGAAGAACTCTTCAC |
| | cvd2019-09-21b | gacctttcgtctctGTCTca**CAAC**ACCAGTGAAGAGTTCTTCACCCTTC |
| GFPN-003 | cvd2019-09-21c | cgatgtacgtctcaCTCG**CC**ATGAGTAAAGGGGAGGAACTCTTTAC |
| | cvd2019-09-21d | gacctttcgtctctGTCTca**CAAC**CCCGGTAAAGAGTTCCTCCCCTTTAC |
| GFPN-004 | cvd2019-09-21e | cgatgtacgtctcaCTCG**CC**ATGTCGAAGGGCGAAGAACTCTTC |
| | cvd2019-09-21f | gacctttcgtctctGTCTca**CAAC**ACCAGTGAAGAGTTCTTCGCCCTTC |
| GFPN-005 | cvd2019-09-21g | cgatgtacgtctcaCTCG**CC**ATGTCTAAGGGTGAGGAGCTCTTC |
| | cvd2019-09-21h | gacctttcgtctctGTCTca**CAAC**TCCCGTGAAGAGCTCCTCACCCTTAG |
| GFPN-006 | cvd2019-09-21i | cgatgtacgtctcaCTCG**CC**ATGTCGAAAGGGGAAGAACTGTTCAC |
| | cvd2019-09-21j | gacctttcgtctctGTCTca**CAAC**GCCGGTGAACAGTTCTTCCCCTTTC |
| GFPN-007 | cvd2019-09-21k | cgatgtacgtctcaCTCG**CC**ATGTCTAAAGGAGAAGAGCTTTTCAC |
| | cvd2019-09-21l | gacctttcgtctctGTCTca**CAAC**CCCAGTGAAAAGCTCTTCTCCTTTAG |
| GFPN-008 | cvd2019-09-21m | cgatgtacgtctcaCTCG**CC**ATGAGTAAGGGTGAGGAATTATTCACG |
| | cvd2019-09-21n | gacctttcgtctctGTCTca**CAAC**GCCCGTGAATAATTCCTCACCCTTAC |
| GFPN-009 | cvd2019-09-22a | cgatgtacgtctcaCTCG**CC**ATGAGTAAAGGGGAAGAACTGTTCAC |
| | cvd2019-09-22b | gacctttcgtctctGTCTca**CAAC**GCCAGTGAACAGTTCTTCCCCTTTAC |
| GFPN-010 | cvd2019-09-22c | cgatgtacgtctcaCTCG**CC**ATGTCTAAGGGTGAGGAGCTCTTC |
| | cvd2019-09-22d | gacctttcgtctctGTCTca**CAAC**TCCTGTGAAGAGCTCCTCACCCTTAG |
| GFPN-011 | cvd2019-09-22e | cgatgtacgtctcaCTCG**CC**ATGAGTAAAGGAGAAGAGTTATTTACTGG |
| | cvd2019-09-22f | gacctttcgtctctGTCTca**CAAC**TCCAGTAAATAACTCTTCTCCTTTAC |
| GFPN-012 | cvd2019-09-22g | cgatgtacgtctcaCTCG**CC**ATGAGTAAGGGAGAAGAGCTGTTC |
| | cvd2019-09-22h | gacctttcgtctctGTCTca**CAAC**TCCAGTGAACAGCTCTTCTCCCTTAC |

| | | |
|---|---|---|
| GFPN-013 | cvd2019-09-22i | cgatgtacgtctcaCTCG**CCATG**TCGAAAGGAGAAGAATTGTTCAC |
| | cvd2019-09-22j | gacctttcgtctctGTCTca**CAAC**GCCCGTGAACAATTCTTCTCCTTTCG |
| GFPN-014 | cvd2019-09-22k | cgatgtacgtctcaCTCG**CCATG**AGCAAAGGAGAAGAATTATTTACTGG |
| | cvd2019-09-22l | gacctttcgtctctGTCTca**CAAC**TCCAGTAAATAATTCTTCTCCTTTGC |
| GFPN-015 | cvd2019-09-22m | cgatgtacgtctcaCTCG**CCATG**AGCAAAGGAGAAGAATTATTTACGG |
| | cvd2019-09-22n | gacctttcgtctctGTCTca**CAAC**TCCCGTAAATAATTCTTCTCCTTTGC |
| GFPN-016 | cvd2019-09-22o | cgatgtacgtctcaCTCG**CCATG**AGCAAAGGGGAAGAATTATTTACAG |
| | cvd2019-09-22p | gacctttcgtctctGTCTca**CAAC**ACCTGTAAATAATTCTTCCCCTTTGC |
| GFPN-017 | cvd2020-03-05a | cgatgtacgtctcaCTCG**CCATG**AGTAAAGGGGAAGAACTCTTTACC |
| | cvd2020-03-05b | gacctttcgtctctGTCTca**CAAC**CCCGGTAAAGAGTTCTTCCCCTTTAC |
| GFPN-018 | cvd2020-03-05c | cgatgtacgtctcaCTCG**CCATG**TCGAAAGGTGAGGAACTATTCACTG |
| | cvd2020-03-05d | gacctttcgtctctGTCTca**CAAC**ACCAGTGAATAGTTCCTCACCTTTC |
| GFPN-019 | cvd2020-03-05e | cgatgtacgtctcaCTCG**CCATG**TCGAAGGGTGAAGAACTGTTCACTG |
| | cvd2020-03-05f | gacctttcgtctctGTCTca**CAAC**ACCAGTGAACAGTTCTTCACCCTTC |
| GFPN-020 | cvd2020-03-05g | cgatgtacgtctcaCTCG**CCATG**TCGAAGGGTGAAGAACTTTTCACTG |
| | cvd2020-03-05h | gacctttcgtctctGTCTca**CAAC**CCCAGTGAAAAGTTCTTCACCCTTC |
| GFPN-021 | cvd2020-03-05i | cgatgtacgtctcaCTCG**CCATG**TCCAAAGGGGAGGAACTCTTTACG |
| | cvd2020-03-05j | gacctttcgtctctGTCTca**CAAC**GCCCGTAAAGAGTTCCTCCCCTTTG |
| GFPN-022 | cvd2020-03-05k | cgatgtacgtctcaCTCG**CCATG**TCCAAAGGTGAAGAGCTTTTCACC |
| | cvd2020-03-05l | gacctttcgtctctGTCTca**CAAC**CCCGGTGAAAAGCTCTTCACCTTTG |
| GFPN-023 | cvd2020-03-05m | cgatgtacgtctcaCTCG**CCATG**TCGAAAGGTGAAGAGCTGTTCAC |
| | cvd2020-03-05n | gacctttcgtctctGTCTca**CAAC**ACCGGTGAACAGCTCTTCACCTTTC |
| GFPN-024 | cvd2020-03-05o | cgatgtacgtctcaCTCG**CCATG**TCGAAAGGTGAGGAACTGTTCAC |
| | cvd2020-03-05p | gacctttcgtctctGTCTca**CAAC**CCCAGTGAACAGTTCCTCACCTTTC |
| GFPN-025 | cvd2020-03-05q | cgatgtacgtctcaCTCG**CCATG**AGTAAGGGGGAGGAGCTCTTCAC |
| | cvd2020-03-05r | gacctttcgtctctGTCTca**CAAC**TCCGGTGAAGAGCTCCTCCCCCTTAC |
| GFPN-026 | cvd2020-03-05s | cgatgtacgtctcaCTCG**CCATG**AGTAAAGGGGAAGAGCTTTTCAC |
| | cvd2020-03-05t | gacctttcgtctctGTCTca**CAAC**CCCGGTGAAAAGCTCTTCCCCTTTAC |
| GFPN-027 | cvd2020-03-06a | cgatgtacgtctcaCTCG**CCATG**AGTAAAGGAGAAGAACTCTTTACCG |

| | | |
|---|---|---|
| | cvd2020-03-06b | gacctttcgtctctGTCTca**CAAC**TCCGGTAAAGAGTTCTTCTCCTTTAC |
| GFPN-028 | cvd2020-03-06c | cgatgtacgtctcaCTCG**CCAT**GAGTAAAGGAGAAGAACTCTTCACC |
| | cvd2020-03-06d | gacctttcgtctctGTCTca**CAAC**ACCGGTGAAGAGTTCTTCTCCTTTAC |
| GFPN-029 | cvd2020-03-06e | cgatgtacgtctcaCTCG**CCAT**GTCAAAGGGGGAAGAACTGTTCAC |
| | cvd2020-03-06f | gacctttcgtctctGTCTca**CAAC**GCCTGTGAACAGTTCTTCCCCCTTTG |
| GFPN-030 | cvd2020-03-06g | cgatgtacgtctcaCTCG**CCAT**GTCGAAAGGCGAGGAACTGTTCAC |
| | cvd2020-03-06h | gacctttcgtctctGTCTca**CAAC**TCCAGTGAACAGTTCCTCGCCTTTC |
| GFPN-031 | cvd2020-03-06i | cgatgtacgtctcaCTCG**CCAT**GAGCAAGGGTGAAGAGTTATTCACTG |
| | cvd2020-03-06j | gacctttcgtctctGTCTca**CAAC**TCCAGTGAATAACTCTTCACCCTTG |
| GFPN-032 | cvd2020-03-06k | cgatgtacgtctcaCTCG**CCAT**GTCTAAAGGTGAAGAACTATTCACAGG |
| | cvd2020-03-06l | gacctttcgtctctGTCTca**CAAC**CCCTGTGAATAGTTCTTCACCTTTAG |
| GFPN-033 | cvd2020-03-06m | cgatgtacgtctcaCTCG**CCAT**GTCTAAAGGTGAGGAGCTCTTCAC |
| | cvd2020-03-06n | gacctttcgtctctGTCTca**CAAC**TCCTGTGAAGAGCTCCTCACCTTTAG |
| GFPN-034 | cvd2020-03-06o | cgatgtacgtctcaCTCG**CCAT**GAGTAAGGGAGAGGAACTGTTCAC |
| | cvd2020-03-06p | gacctttcgtctctGTCTca**CAAC**CCCTGTGAACAGTTCCTCTCCCTTAC |
| GFPN-035 | cvd2020-03-06q | cgatgtacgtctcaCTCG**CCAT**GTCGAAAGGGGAAGAATTGTTCAC |
| | cvd2020-03-06r | gacctttcgtctctGTCTca**CAAC**TCCAGTGAACAATTCTTCCCCTTTCG |
| GFPN-036 | cvd2020-03-06s | cgatgtacgtctcaCTCG**CCAT**GAGTAAGGGGGAGGAGCTGTTC |
| | cvd2020-03-06t | gacctttcgtctctGTCTca**CAAC**TCCTGTGAACAGCTCCTCCCCCTTAC |
| GFPN-037 | cvd2020-03-07a | cgatgtacgtctcaCTCG**CCAT**GAGTAAGGGAGAGGAATTGTTCAC |
| | cvd2020-03-07b | gacctttcgtctctGTCTca**CAAC**ACCCGTGAACAATTCCTCTCCCTTAC |
| GFPN-038 | cvd2020-03-07c | cgatgtacgtctcaCTCG**CCAT**GAGTAAGGGAGAGGAACTTTTCAC |
| | cvd2020-03-07d | gacctttcgtctctGTCTca**CAAC**TCCCGTGAAAAGTTCCTCTCCCTTAC |
| GFPN-039 | cvd2020-03-07e | cgatgtacgtctcaCTCG**CCAT**GAGTAAAGGAGAGGAGCTTTTCACAG |
| | cvd2020-03-07f | gacctttcgtctctGTCTca**CAAC**TCCTGTGAAAAGCTCCTCTCCTTTAC |
| GFPN-040 | cvd2020-03-07g | cgatgtacgtctcaCTCG**CCAT**GAGCAAAGGAGAAGAGTTATTTACAGG |
| | cvd2020-03-07h | gacctttcgtctctGTCTca**CAAC**CCCTGTAAATAACTCTTCTCCTTTGC |
| GFPN-041 | cvd2020-03-07i | cgatgtacgtctcaCTCG**CCAT**GAGCAAAGGAGAGGAATTATTTACG |
| | cvd2020-03-07j | gacctttcgtctctGTCTca**CAAC**GCCCGTAAATAATTCCTCTCCTTTGC |

| | | |
|---|---|---|
| GFPN-042 | cvd2020-05-15a | cgatgtacgtctcaCTCG**CCAT**GAGTAAAGGGGAGGAACTCTTTACTG |
| | cvd2020-05-15b | gacctttcgtctctGTCTca**CAAC**ACCAGTAAAGAGTTCCTCCCCTTTAC |
| GFPN-043 | cvd2020-05-15c | cgatgtacgtctcaCTCG**CCAT**GTCGAAAGGTGAAGAACTTTTCACTG |
| | cvd2020-05-15d | gacctttcgtctctGTCTca**CAAC**ACCAGTGAAAAGTTCTTCACCTTTCG |
| GFPN-044 | cvd2020-05-15e | cgatgtacgtctcaCTCG**CCAT**GAGCAAGGGAGAAGAGCTGTTCACTG |
| | cvd2020-05-15f | gacctttcgtctctGTCTca**CAAC**GCCAGTGAACAGCTCTTCTCCC |
| GFPN-045 | cvd2020-05-15g | cgatgtacgtctcaCTCG**CCAT**GAGTAAGGGTGAGGAGTTATTCACG |
| | cvd2020-05-15h | gacctttcgtctctGTCTca**CAAC**GCCCGTGAATAACTCCTCACCCTTAC |
| GFPN-046 | cvd2020-05-15i | cgatgtacgtctcaCTCG**CCAT**GTCTAAAGGAGAAGAACTCTTCACAGG |
| | cvd2020-05-15j | gacctttcgtctctGTCTca**CAAC**CCCTGTGAAGAGTTCTTCTCCTTTAG |
| GFPN-047 | cvd2020-05-15k | cgatgtacgtctcaCTCG**CCAT**GTCCAAAGGAGAAGAACTATTCACC |
| | cvd2020-05-15l | gacctttcgtctctGTCTca**CAAC**TCCGGTGAATAGTTCTTCTCCTTTGG |
| GFPN-048 | cvd2020-05-15m | cgatgtacgtctcaCTCG**CCAT**GAGCAAAGGAGAAGAACTATTCACG |
| | cvd2020-05-15n | gacctttcgtctctGTCTca**CAAC**TCCCGTGAATAGTTCTTCTCCTTTGC |
| GFPN-049 | cvd2020-05-15o | cgatgtacgtctcaCTCG**CCAT**GTCGAAGGGAGAAGAATTATTTACGG |
| | cvd2020-05-15p | gacctttcgtctctGTCTca**CAAC**TCCCGTAAATAATTCTTCTCCCTTCG |

**Table S5. Oligonucleotide primer pairs for constructing TIsigner variants of luciferase.** The sequences of each forward and reverse primer pair used for constructing each of the luciferase TIsigner variants is shown. The start codon in each of the forward primers is shaded yellow. BsmBI recognition sites (used for Golden Gate assembly into the MIDAS pML1 vector) are underlined. The MIDAS prefix [CCAT] and suffix [AGGA] (reverse-complement = TCCT) for the RLucN modules are highlighted in bold blue and red, respectively.

| RLucN TIsigner ID | Oligonucleotide Primer Pair | Primer Sequences (5' to 3') |
|---|---|---|
| RLucN-TI-002 | cvd2019-06-14c | cgatgtacgtctcaCTCG**CCAT**GACATCAAAAGTATACGACCCAGAG |
| | cvd2019-06-14d | gacctttcgtctctGTCTca**TCCT**CTGCTCTGGGTCGTATACTTTTGATG |
| RLucN-TI-003 | cvd2019-06-14e | cgatgtacgtctcaCTCG**CCAT**GACAAGTAAAGTTTATGACCCAGAGC |
| | cvd2019-06-14f | gacctttcgtctctGTCTca**TCCT**CTGCTCTGGGTCATAAACTTTACTTG |
| RLucN-TI-004 | cvd2019-06-14g | cgatgtacgtctcaCTCG**CCAT**GACCAGCAAAGTTTATGACCCAGAG |
| | cvd2019-06-14h | gacctttcgtctctGTCTca**TCCT**CTGCTCTGGGTCATAAACTTTGCTG |
| RLucN-TI-005 | cvd2019-06-14i | cgatgtacgtctcaCTCG**CCAT**GACAAGCAAAGTTTATGACCCAGAGC |
| | cvd2019-06-14j | gacctttcgtctctGTCTca**TCCT**CTGCTCTGGGTCATAAACTTTGC |
| RLucN-TI-006 | cvd2019-06-14k | cgatgtacgtctcaCTCG**CCAT**GACTTCGAAAGTTTATGATCCAGAACAG |
| | cvd2019-06-14l | gacctttcgtctctGTCTca**TCCT**CTGTTCTGGATCATAAACTTTCGAAG |
| RLucN-TI-007 | cvd2019-06-14m | cgatgtacgtctcaCTCG**CCAT**GACATCAAAAGTTTATGATCCAGAACAAAG |
| | cvd2019-06-14n | gacctttcgtctctGTCTca**TCCT**TTGTTCTGGATCATAAACTTTTGATGTC |
| RLucN-TI-008 | cvd2019-06-14o | cgatgtacgtctcaCTCG**CCAT**GACGTCGAAAGTTTACGATCCAG |
| | cvd2019-06-14p | gacctttcgtctctGTCTca**TCCT**TTGTTCTGGATCGTAAACTTTCGACG |
| RLucN-TI-009 | cvd2019-06-14q | cgatgtacgtctcaCTCG**CCAT**GACATCGAAAGTTTACGATCCAGAAC |
| | cvd2019-06-14r | gacctttcgtctctGTCTca**TCCT**TTGTTCTGGATCGTAAACTTTCGATG |
| RLucN-TI-010 | cvd2019-06-14s | cgatgtacgtctcaCTCG**CCAT**GACCTCGAAAGTTTATGACCCAGAAC |
| | cvd2019-06-14t | gacctttcgtctctGTCTca**TCCT**TTGTTCTGGGTCATAAACTTTCGAG |
